# Bilateral cellular flows display asymmetry prior to left-right organizer formation in amniote gastrulation

**DOI:** 10.1101/2024.04.21.590437

**Authors:** Rieko Asai, Shubham Sinha, Vivek N. Prakash, Takashi Mikawa

## Abstract

A bilateral body plan is predominant throughout the animal kingdom. Bilaterality of amniote embryos becomes recognizable as midline morphogenesis begins at gastrulation, bisecting an embryonic field into the left and right sides, and left-right asymmetry patterning follows. While a series of laterality genes expressed after the left-right compartmentalization has been extensively studied, the laterality patterning prior to and at the initiation of midline morphogenesis has remained unclear. Here, through a biophysical quantification in a high spatial and temporal resolution, applied to a chick model system, we show that a large-scale bilateral counter-rotating cellular flow, termed as ‘polonaise movements’, display left-right asymmetries in early gastrulation. This cell movement starts prior to the formation of the primitive streak (the earliest midline structure) and the subsequent appearance of Hensen’s node (the left-right organizer). The cellular flow speed and vorticity unravel the location and timing of the left-right asymmetries. The bilateral flows displayed a Right dominance after six hours since the start of cell movements. Mitotic arrest that diminishes primitive streak formation resulted in changes in the bilateral flow pattern, but the Right dominance persisted. Our data indicate that the left-right asymmetry in amniote gastrula becomes detectable earlier than suggested by current models, which assume that the asymmetric regulation of the laterality signals at the node leads to the left-right patterning. More broadly, our results suggest that physical processes can play an unexpected but significant role in influencing left-right laterality during embryonic development.

**Significance Statement:** Bilaterians are defined by a bilaterally symmetrical body plan. Vertebrates exhibit external bilateral symmetry but display left-right (LR) asymmetry in their internal organs. Amniote embryos switch the patterning of internal organs from bilateral symmetry to LR-asymmetry. Using chick embryos as a model system, here we examined the initiation of LR symmetry breaking. Our biophysical approaches to quantify cellular flows inferred that LR symmetry breaking occurs before the formation of Hensen’s node, a LR organizer, which serves as a signaling center for LR patterning-gene programs. Our work demonstrates that quantitative biophysical parameters can help unravel the initiation of LR symmetry breaking, suggesting an involvement of physical mechanisms in this critical biological patterning process.

## Introduction

Most animals are classified in the *Bilateria*, possessing a morphological symmetry between the left-right (LR) sides conferred by the midline. Vertebrates present a complex bilateral body plan, displaying bilateral symmetry externally but with asymmetry in the internal organs (1-3). LR patterning and morphogenesis have been extensively studied for events post-formation of Hensen’s node in amniotes (Kupffer’s vesicle in fish)(4, 5), which induces the asymmetric expression and regulation of the LR regulatory genes (6, 7).

In amniote embryos, the bilateral body plan becomes recognizable at gastrulation during midline morphogenesis, compartmentalizing the embryonic field into the left and right sides (7, 8). The primitive streak (PS) is the earliest midline recognizable landmark and serves as a signaling center of gastrulation (9, 10). Prior to and during PS extension, a counter-rotating cellular flow, termed as ‘polonaise movements’, appears at both LR sides along the midline axis (11-17). This bilateral rotating cellular flow continues until ingression to generate germ layers starts, at the midline through the PS. At this point, the cellular flow shifts from a rotation pattern to a lateral-to-medial pattern toward the midline or PS (18). Previously, we have shown that the lateral-to-medial movements of the epiblast demonstrate ipsilaterality, or side-identity, along the PS (19). Furthermore, a LR-asymmetric membrane potential within the extending PS has been reported (20). These studies hint at a possibility that LR laterality might arise even before the genetically programmed LR patterning is established.

Here, we examine the earliest timing of LR asymmetry patterning in amniote gastrulation. Utilizing computational analyses based on physics of fluids (21), we examined the cellular flows associated with the polonaise movements during early gastrulation in the chick embryo. Visual mapping of biophysical parameters of cellular flows over time revealed that the cellular flows on the right side were dominant than in the left during PS formation. To test the role of midline structure, PS formation was diminished through mitotic arrest. We show that the resulting embryos displayed a shortened duration of bilateral cellular flows and ended with total right-side dominance of the cellular flows. These data suggest that the laterality program regulating the right-side dominance of the cellular flows is already set up prior to both maturation of the midline and LR asymmetric expression of the well-known laterality genes at Hensen’s node, and it does not depend on cell division.

## Results

### Bilateral flows during early midline morphogenesis in chick gastrulation

To survey the timing of initiation and patterning of the bilateral cellular flow, we performed live-imaging experiments during the early chick gastrulation process, from pre-streak stage until HH3 [Hamburger and Hamilton staging (22)] (Figure 1 a, b, d) (Methods). To quantitatively analyze the cellular flow during early midline morphogenesis or PS formation, the image sequences of fluorescently tagged cells (Methods) were post-processed using the Particle Image Velocimetry (PIV) method (21, 23). PIV is a combined experimental and computational technique from fluid mechanics that is used to quantify time-varying fluid velocity fields by analyzing time-lapse datasets with moving tracer particles (Methods). A snapshot of the flow velocity vectors is shown in Figure 1c, indicating the presence of the well-known polonaise movements in our experiments (Movie 1).

**Figure 1.**
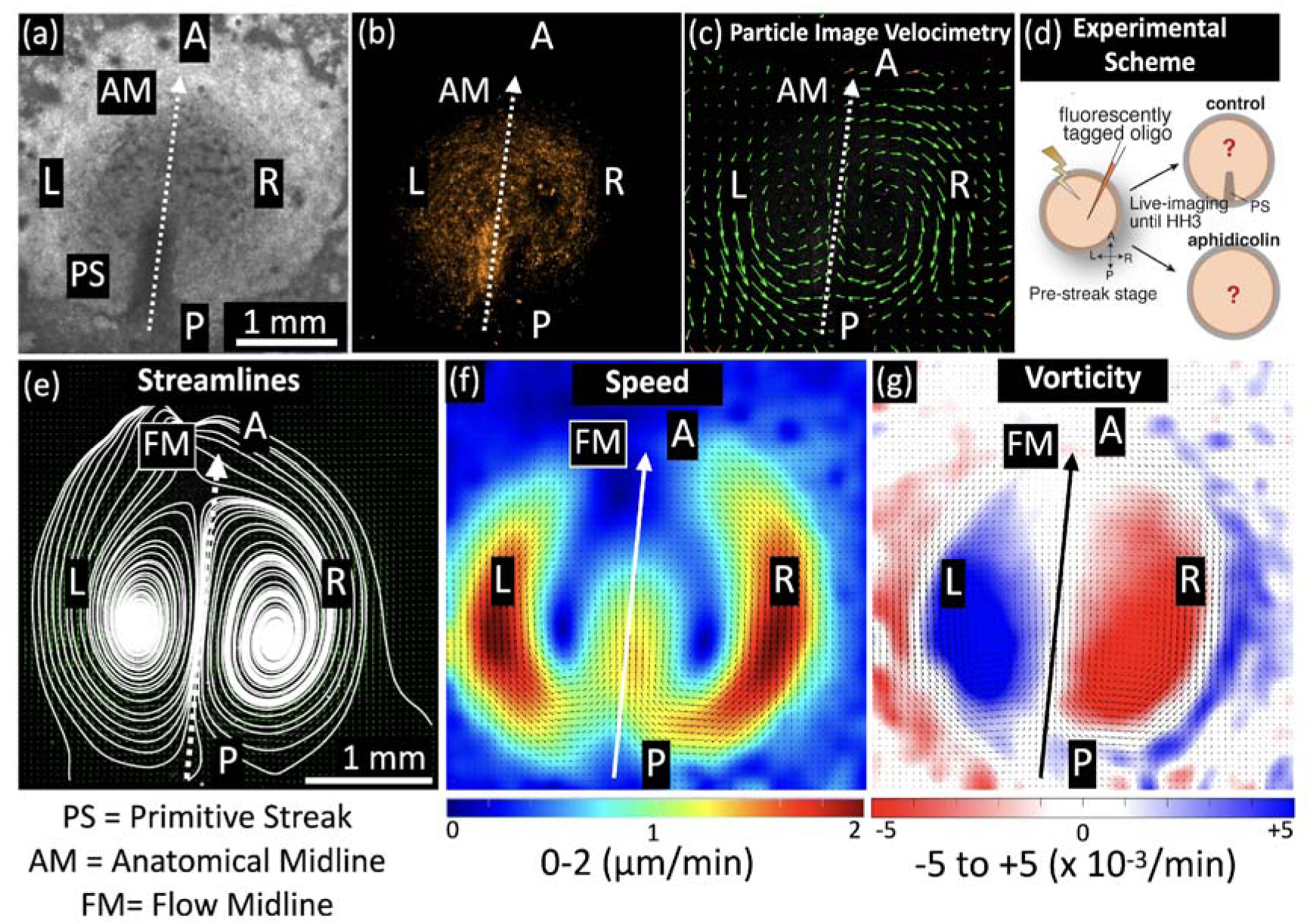
Cellular flows during early chick embryo development. **Upper panel:** (a) Brightfield image of chicken embryo gastrulation showing development of the primitive Streak (PS) at the midline (at stage HH3). The Anatomical Midline (AM) defined by the PS is shown in dotted lines, and the Anterior (A), Posterior (P), Left (L) and Right (R) regions are also labeled. The images shown in (a-c) correspond to the time point of 10 hours post initiation of cellular motion in control sample 1 (see Table 1). (b) Fluorescence image of the embryo revealing cells tagged with fluorescent markers. (c) Results from Particle Image Velocimetry (PIV) analysis (Methods), the green arrows represent instantaneous velocity vector fields. (d) A schematic showing the experimental scheme (Methods). **Lower panel:** (e-g) Time-averaged (over 10 hours) results from PIV analysis. FM is the Flow Midline (see Figure 4). (e) Streamlines reveal bilateral cellular flow patterns with counter-rotating vortices in both L and R regions. (f) Cellular speeds: heat map indicates lowest speeds in blue color and highest speeds in red color. (g) Vorticity (measurement of rotation): Red colors indicate clockwise rotation, and blue colors indicate counterclockwise rotation directions. Both speed and vorticity plots show slightly higher magnitudes on the Right side, indicating asymmetry in the bilateral flows (f, g). The length scale bar shown in (a) is the same for panels (a-c), and length scale bar in (e) is the same for panels (e-g).

**Table 1.**
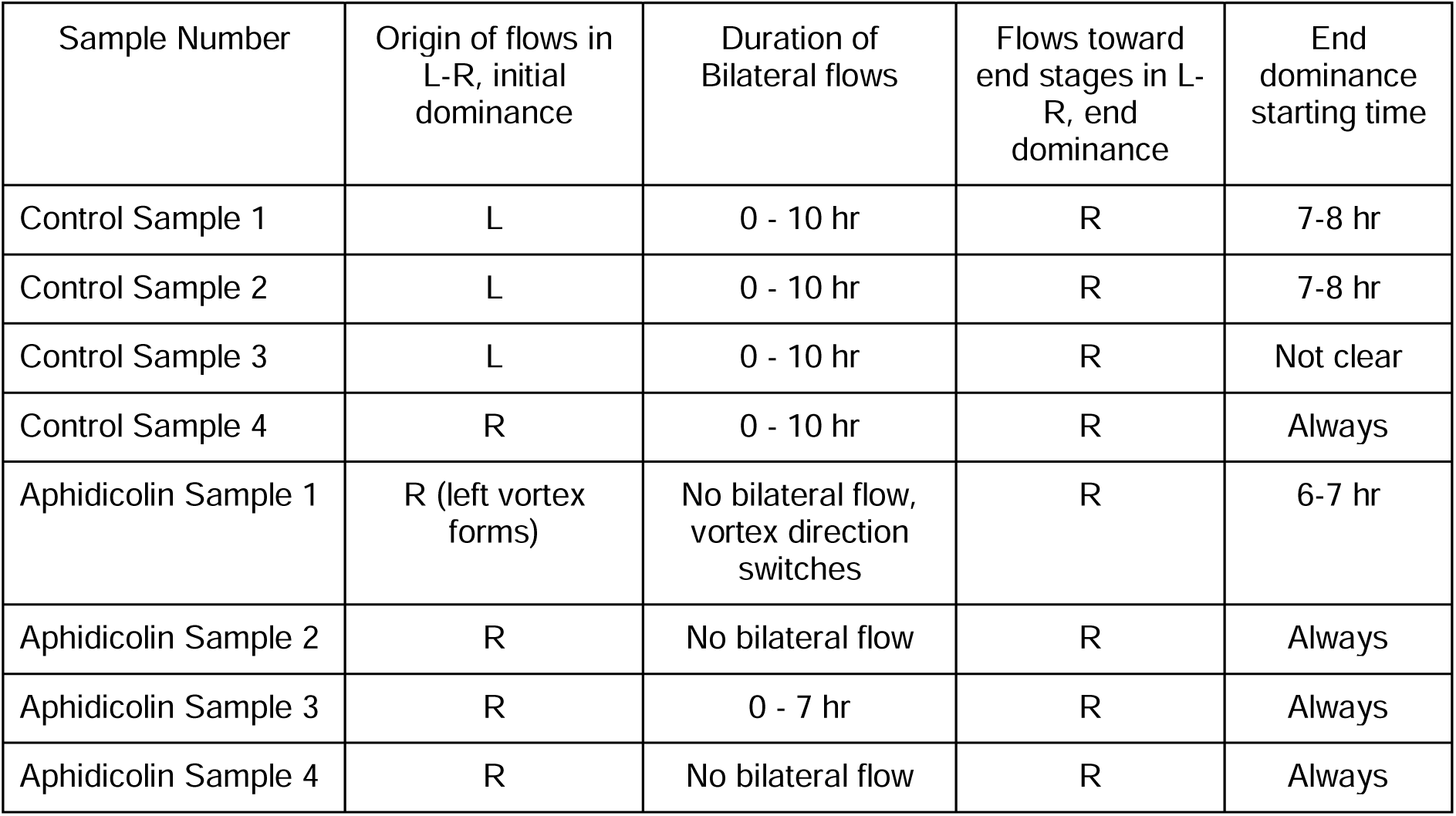
Features of bilateral flows in control versus aphidicolin-treated embryos.

To obtain a comprehensive pattern of the bilateral cellular flows, time-averaged data of the flow fields were generated for the entire imaging duration (10 hours) (Figure 1 e-g) (Methods). The time-averaged flow streamlines revealed the intricate structure of two counter-rotating vortices that characterize the bilateral cellular flows (Figure 1e). The time-averaged speed calculations showed that the fastest cellular flows were in the peripheral regions of the vortices, on both L and R sides (Figure 1f). The bilateral flows also had a local rotational component that was quantified using a metric known as vorticity. We extracted the time-averaged vorticity information, and the two counter-rotating vortices were visible in L and R sides (blue- and red-colored regions, respectively) (Figure 1g). The darker colored regions in L and R (blue- and red-colored, respectively) indicate highest vorticity magnitudes, corresponding to the location of the bilateral vortices (Figure 1g). The vorticity values considered for further quantification were only those lying inside the embryonic region (Figure S1).

Strikingly, the full time-averaged analysis provided higher resolution data for spatiotemporal patterning of the bilateral cellular flows. Looking closely at the speed plot (Figure 1f), the regions with higher speed on the R side (yellow and red regions) qualitatively occupy a slightly larger area than on the L side and quantification of this area is carried out in sections that follow. Similarly, the vorticity regions in R (red) qualitatively occupied a larger area than the L (blue) (Figure 1g). These time-averaged results suggest that the bilateral flows may not be fully symmetric. A LR asymmetry, more specifically a R dominance, became evident (Figure 1 f, g).

### Temporal changes in bilateral flows during early midline morphogenesis in chick gastrulation

To study the time-varying bilateral cellular flow patterns in the L and R, we performed the PIV averaging by generating hourly time-averaged plots for the entire 10 hours of imaging (Figure 2). The cellular flows were accompanied by formation of the two counter-rotating vortices (see streamlines, Figure 2). In the initial hours (0-2 hours), these vortices were not fully developed. Whereas, at later time points, they formed two fully developed counter-rotating vortices (4 hours onwards) (Figure 2). The speed plots showed that the cellular flows initiated along the peripheral regions (0-2 hours) and merged at the posterior end. The flows then proceeded from posterior (P) to anterior (A) along the Flow Midline (FM) axis (3 hours onwards) (Figure 2). In the next 4 hours (3-7 hours), the L and R sides in the speed plots presented high symmetry. At later times (7-8 hours), the flow on the R side dominates with a larger area containing higher speeds, and this R domination continues till the end. The streamlines also showed that at later times the R vortex becomes larger in size than the L vortex. The vorticity calculations also showed initial unstable transients (0-2 hours) but stabilized after a few hours (Figure 2). As the bilateral flows proceeded, the L (blue) and R (red) side vortices grew in size to form stable counter-rotating vortices. At a later timepoint (6 hours onwards), the R (red) vortex was getting slightly larger in area compared to the L (blue) vortex. This R side vorticity dominance further increased and persisted throughout this window (9-10 hours).

**Figure 2.**
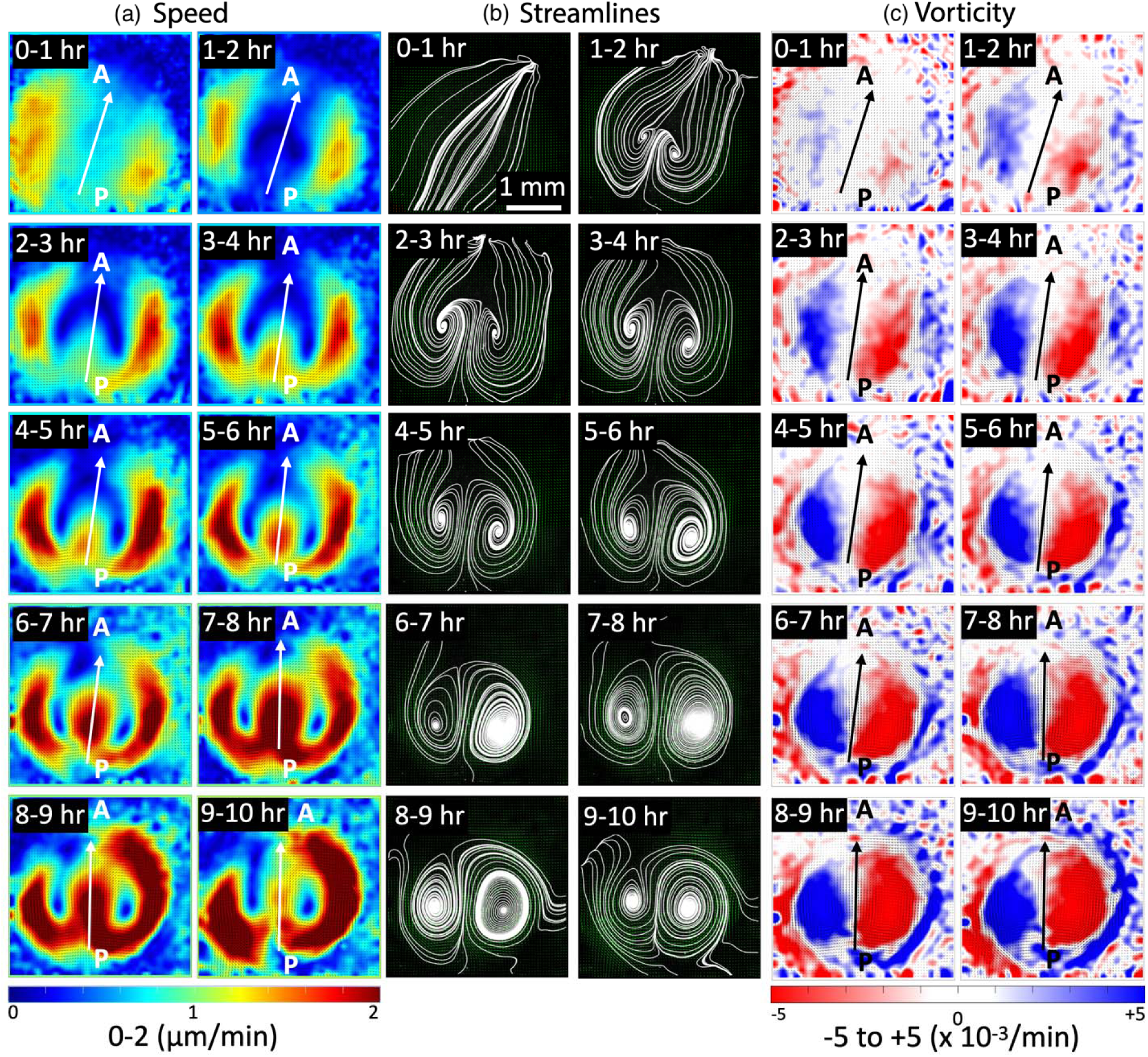
Temporal progression of bilateral flows during early chick embryo development. Short time-averaged (1 hour) sequences of (a) speed, (b) streamlines, and (c) vorticity from PIV analysis of control sample 1 (see Table 1). (a) Speed: cellular flows show initial speed transients (0-2 hours), which later stabilize and lead to symmetric L-R speed patterns (2-7 hours). This symmetry in flow speed is lost at the later stages (7 hours onwards) and cellular speeds become R dominant. (b) Streamlines: Cellular flows initiate only at the sides (0-1 hours), and then progress in the midline region displayed by the bilateral transients (1-3 hours). Later these bilateral flows form two stable counter-rotating vortices (3 hours onwards). (c) Vorticity: Initial flow transients (0-2 hours) stabilize to form symmetric bilateral flows (2-7 hours) with well-defined counter-rotating vortices (blue-red colors). Following this (7 hours onwards), the L-R symmetry is lost, with the R vortex being dominant. The arrows represent the Flow Midlines (FM) and indicate the direction of flows at that location (Methods). The length scale bar shown in the first streamline panel (0-1 hour) is the same for all panels.

In another dataset (control sample 2), the bilateral flows initially began with a L dominance and at the end transition to R dominant flows (Figure S2) as seen with other samples. In this study, we define the ‘dominance’ of the cellular flow to explicitly describe that flow strength is larger on one side compared to the other. Table 1 lists the origin and the end dominance of flow in all the four control samples in this study. All the cases exhibited R dominant flow at later times. In summary, during early midline morphogenesis, i.e. PS formation during gastrulation in chick embryos, there was a temporal flow transition from L to R dominance pointing that cellular flows exhibit LR asymmetry.

To test whether known laterality genes are detectable at and during LR asymmetry of cellular flow as early midline morphogenesis proceeds, the expression patterns of well-established LR regulatory genes (e.g., *Shh*, *Lefty*, and *Nodal*) were examined during PS extension by whole-mount in situ hybridization (WISH) (Figure 3, Figure S3, Methods). At pre-streak stage X [Eyal-Giladi and Kovchav staging (24)], the expression of *Brachyury* (a PS and early mesodermal cell marker) was not detectable by NBT/BCIP staining (Figure 3, Figure S3). Upon elongation along the midline, PS cells became distinguishable from other embryonic cells as the area of *Brachyury-positive* (a PS and early mesodermal cell marker) and *Sox3-negative* (an epiblast marker) expression (Figure S3). The PS appears from the posterior marginal zone at HH2 (Figure S3) and anteriorly extends along the midline (Figure S3), until the Hensen’s node, which is called as “the LR-organizer” in bird embryos, formed at the anterior tip of the PS at HH4 (22). The NBT/BCIP staining showed that the *Brachyury*-expression within the PS was detectable within the extending PS, while the LR asymmetric expression pattern of the LR regulatory genes became remarkable at Hensen’s node at or after HH4 (Figure S3), consistent with previous reports (25-27). It must be noted that the classical WISH methods, which have been commonly used in previous (25-27) and the present studies (Methods), have a technical limitation of lower sensitivity than the latest RNA detection techniques, such as Hybridization Chain Reaction (HCR) (28). The bilateral cellular flows display qualitatively symmetric flow at HH2 but exhibit asymmetry with a R dominance at HH3 (Figure 3c). Thus, within the technical limitations of the WISH method, LR asymmetries in the cellular flows were detected prior to the expression of well-established LR regulatory genes (e.g., *Shh*, *Lefty*, and *Nodal*) in distinct LR asymmetric patterns at Hensen’s node.

**Figure 3.**
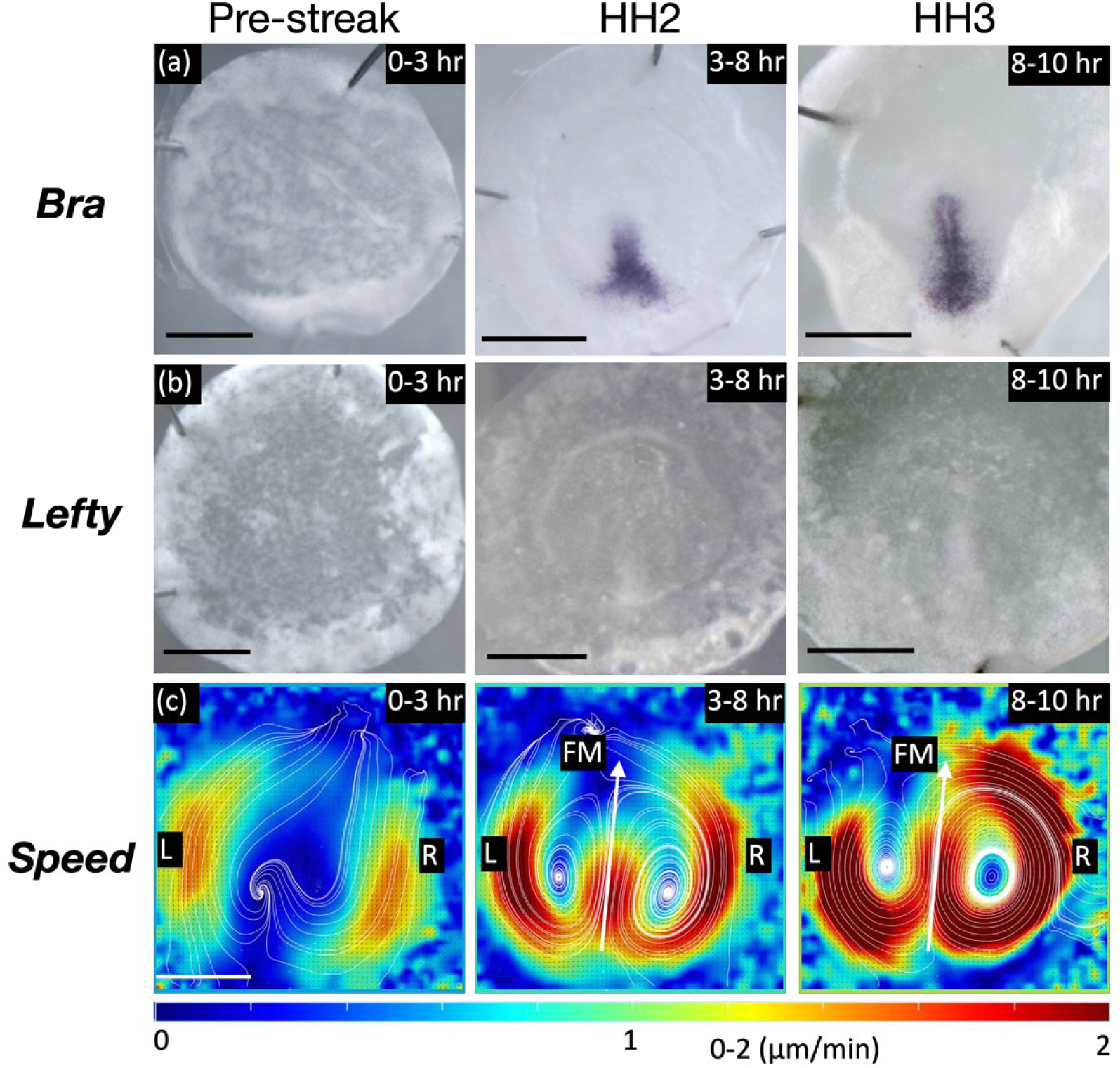
Expression analysis of Lefty (a laterality gene) and the cellular flow. (a, b) Pre-strea stage to HH3 whole-mount in situ hybridization of *Brachyury* (a), Lefty (b). (c) Heat map of time-averaged speed. Scale bars, 1mm.

### Biophysical quantification of bilateral flows during early midline morphogenesis

To quantify the degree of LR asymmetry, the data were analyzed on the L and R sides separately along a midline axis. Three possible midline axes were defined: Anatomical Midline (AM), Biophysical Midline (BM) and Flow Midline (FM). The AM is the line passing through the middle of the PS, and BM is the perpendicular bisector to the line connecting the bilateral vortex centers. The FM is the line parallel to the cellular flow direction in between the bilateral vortices (Methods) (Figure 4a, Figure S4). If the bilateral vortices were symmetric, these three midlines would coincide with each other. To test this hypothesis, we quantified them over time and measured their angular variation with respect to the vertical (Figure 4b, Figure S4). The angular separation between BM and FM was larger at the initiation of the bilateral flow but reduced when the vortices were fully developed (Figure 4c). Further, the three midline axes were not coincident with each other throughout the time. These results support the model that the bilateral cellular flows are asymmetric.

**Figure 4.**
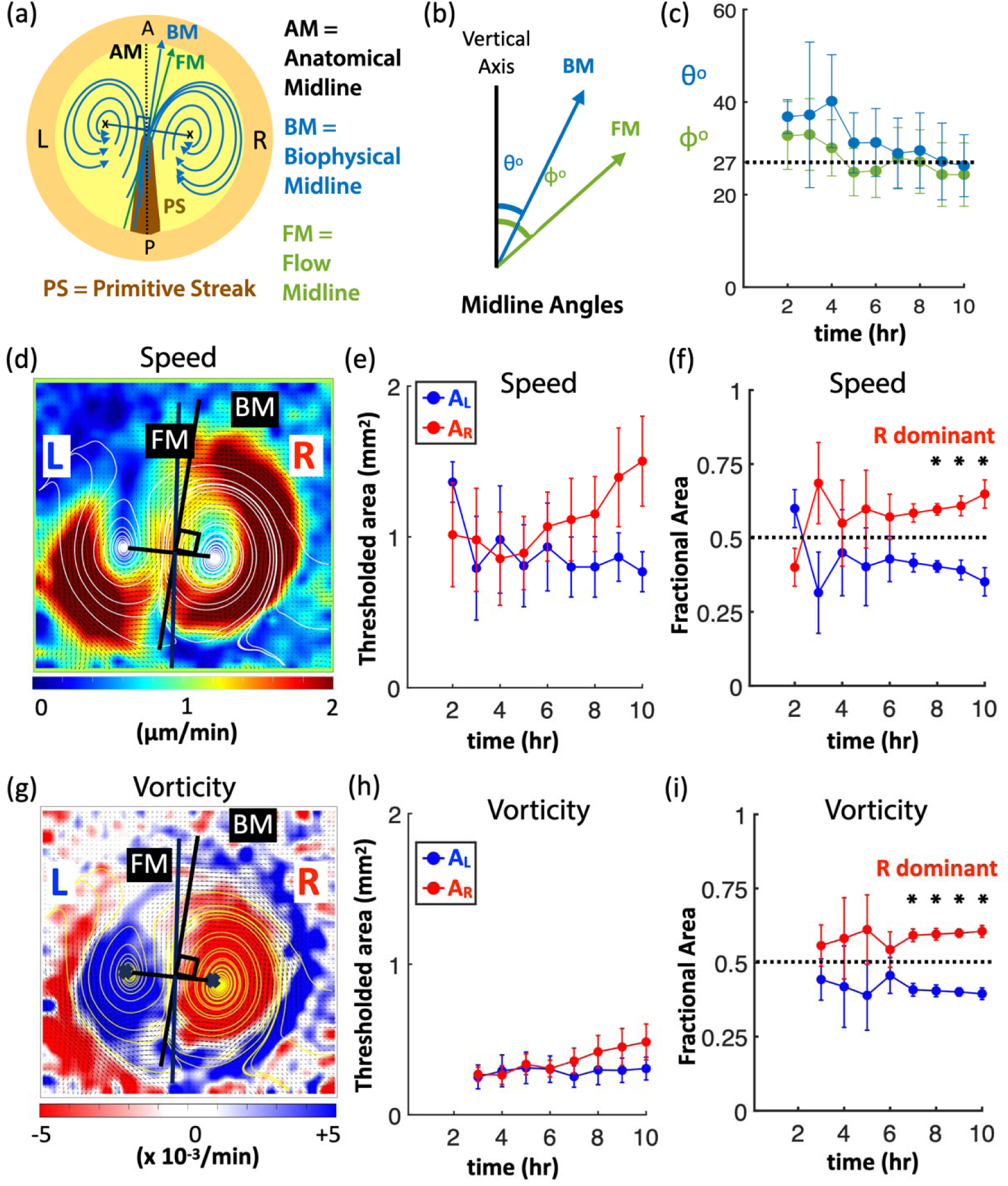
Biophysical quantification of bilateral flows over time. (a) Scheme representation of bilateral flows accompanied by primitive streak (PS) formation. The three distinct midline axes are shown: AM, BM, and FM (supplementary information). (b) The angle subtended by FM and BM to the vertical axis. (c) The averaged (N = 4) mean angles of BM and FM for the control samples. The dotted black line shows the constant angle maintained by AM with the vertical axis. (d) Heat map showing cell flow speeds with the L and R regions of the embryo separated by the FM. (e) The averaged mean (N = 4) of thresholded area of speeds in L (A_L_) and R (A_R_) regions of control samples. (f) The averaged mean (N = 4) of fractional area in L and R regions, corresponding to A_L_/(A_L_+A_R_) and A_R_/(A_L_+A_R_) respectively. (g) Heat map showing the vorticity in L and R regions (blue and red respectively). (h) The averaged mean (N = 4) of thresholded area of vorticity in L (A_L_) and R (A_R_) regions. (i) The averaged mean (N = 4) vorticit fractional area in L and R regions. In (f) and (i), * represents the p-value (0.03) obtained from a non-parametric Wilcoxon test (supplementary information). In (e), (f), (h) and (i), the results correspond to a 50% threshold (supplementary information). The error bars in all plots represents the standard error.

In addition to the midline axes, the positional variations in the L and R vortices were detected over time. This positional variation was quantified by measuring two parameters: positions of the vortex centers (Figure S5) and the distance between the two centers over time (Figure S6). The X and Y locations of the two vortex centers were tracked over the 10-hour duration (Figure S5). The origin (reference point) was chosen to be outside the embryonic disc. The results did not show any significant shift in the X or Y displacements of the vortex centers (N = 4 control samples). Hence, this analysis demonstrated that there was no systematic drift in any direction due to the embryonic disc or the culture system. The average distance between the vortex centers (N = 4) showed larger fluctuations only during the initial hours when the flow was unstable, and later became smaller showing steady fluctuations when the flow stabilized (Figure S6b, c).

In the analyses that follow, we only retained the cellular bilateral flows inside the circular embryonic region. The extra-embryonic material surrounding the embryonic disc was stationary, so our quantification methods gave rise to results that were noise or artifacts, which were discarded (Figure S1). The cellular flow speed and vorticity in the L and R regions showed R dominance (Figures 1 and 2). In addition, the thresholded area of cellular flows for both speed and vorticity increased as the embryos developed. These areas on the L and R regions are denoted as A_L_ and A_R_ respectively. The time evolution of A_L_ and A_R_ for speed and vorticity for all samples with three threshold values (30%, 50%, 70%, Figure S7) are shown in Figures S8 and S9, and the averaged A_L_ and A_R_ for speed and vorticity (N = 4 samples) are shown in Figure S10. Figures 4(e) and (h) show the hourly averaged mean (N= 4 samples) A_L_ and A_R_ of speed and vorticity in L (A_L_) and R (A_R_) regions and their time evolution (50% threshold value is chosen to be the most representative). For both speed and vorticity, we observed that initially (up to 6 hours) A_L_ and A_R_ have similar values. Whereas, at later time points (after 6 hours), A_R_ dominates with larger values (Figures 4e, h). This dominance of A_R_ further increases at the end time points.

The embryonic size variations across different samples were considered by calculating the fractional areas of A_L_ and A_R_ regions, i.e. A_L_/(A_L_+A_R_) and A_R_/(A_L_+A_R_) respectively. Figure 4(f, i) shows the averaged mean (N=4 samples) fractional area for speed and vorticity regions respectively and its time evolution. A value of 0.5 fractional area represents the line of symmetry (represented by the dotted line in Figure 4(f, i)). Any deviation of fractional area value from 0.5 represents asymmetricity. In the beginning stages (up to the initial 6 hours), the L and R fractional areas are closer (symmetric). But at later time points (after 6 hours), the fractional area values separate out with R being dominant (>0.5). The fractional areas showed significant statistical differences in L and R beyond 6 hours, with p-values of 0.03 obtained from a non-parametric Wilcoxon test (see Methods, Supplementary Information). Hence, the bilateral cellular flows showed a clear R dominance at the end stages.

### Bilateral flows in cell-division inhibited embryos during early midline morphogenesis

The above results demonstrated R dominance of the bilateral cellular flows during PS extension. Our recent work has shown that the bilateral cellular flow initiates with diminished PS formation under mitotic arrest (i.e. inhibition of cell division) (29). However, it remains unclear whether proper PS formation was required for R dominance of the bilateral cellular flows. To test the question, we performed live-imaging of the early chick gastrulation process under mitotic arrest caused by aphidicolin (a DNA polymerase inhibitor) (Figure 5, Figure S11, Movie 2). When cell division was inhibited, the PS structure was significantly diminished, consistent with our recent study (Figure S11) (Methods).

**Figure 5.**
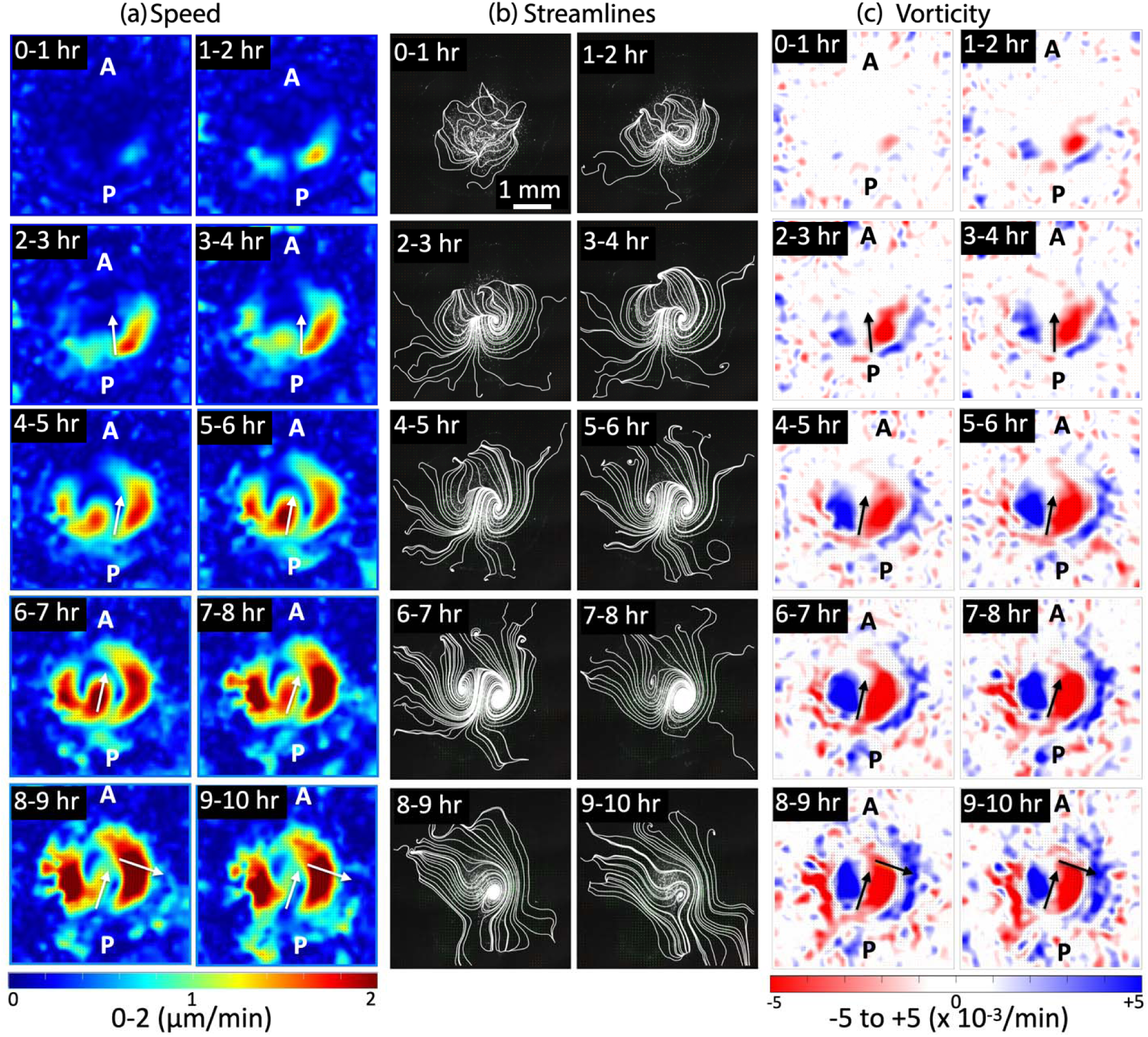
Time evolution of flows in cell-division inhibited chick embryos. Short time-averaged (1 hour) sequences of (a) speeds, (b) streamlines, and (c) vorticity from PIV analysis of aphidicolin sample 3 (see Table 1). (a) Speed: Cellular flows are initially transient and first start on the R side (0-2 hours). These transients stabilize without any L-R symmetry with a dominant R side pattern until the end (2-10 hours). (b) Streamlines: The initial cellular flow transients are random (0-2 hours), but following these bilateral flows develop (2-7 hours). Subsequently, the flows lose L-R symmetry and turn into a single vortex on the R side (7-10 hours). (c) Vorticity: initial flow transients (0-2 hours) develop to form bilateral flows (2-7 hours) with well-defined counter rotating vortices (blue-red colors). After this (7 hours onwards) even though there is only a R vortex present (red), the blue color indicates the counterclockwise rotational regions that still persist (but do not form a closed vortex) (Methods). The short arrows serve as a guide to indicate direction of flows at that location, not representing the midlines (Methods). The length scale bar shown in the first streamline panel (0-1 hour) is the same for all panels.

Under mitotic arrest, the cellular flow patterns in the PS-diminished embryos displayed different patterns from those seen in the control (Figure 5, Movie 2). The flows started with a side dominance (frequently R) (Table 1) forming a bilateral flow briefly (Figure 5). This bilateral flow later merged into one single R vortex. The initial presence of this bilateral flow had already been lost in some cases (Figure S12) (Table 1). However, in all cases (N = 4), the flows merged to form a single vortex with a R dominance at later stages. The temporal evolution of the speeds also showed that the cellular flows were always R-dominant (Figure 5). Similarly, for the vorticity as well, a R dominance was detectable from the initial stages (clockwise rotating vortex region in red color). The blue-colored regions in the vorticity plots correspond to curved flow regions that have counterclockwise rotations, but a closed vortex did not form (confirmed from the corresponding streamlines) (Figure 5). In these embryos that were under mitotic arrest, PS formation was diminished (Figure S11), indicating that the AM was not defined in these embryos. Further, consistent BM and FM were not defined, because the bilateral flows did not continue throughout this time window (Figure 5, Movie 2). While these technical limitations made it difficult to carry out further biophysical quantification, the above PIV analyses clearly identified that R dominance of the bilateral cellular flows persisted under mitotic arrest.

### Comparison of bilateral flows in control versus cell-division inhibited embryos during early midline morphogenesis

The cellular flows in the control and cell-division inhibited embryos are compared and summarized here. Since biophysical quantification indicated that the degree of LR asymmetry of the cellular flow shifted at approximately 5 hours (Figure 4), we calculated the half-time averages (5 hours) of the datasets to account for these temporal flow patterns and variations. These half-time averages capture the flow dynamics during the first half (0-5 hours) and the second half (5-10 hours) of the early midline morphogenesis process in both the control and cell-division inhibited cases, as shown in Figure 6.

**Figure 6.**
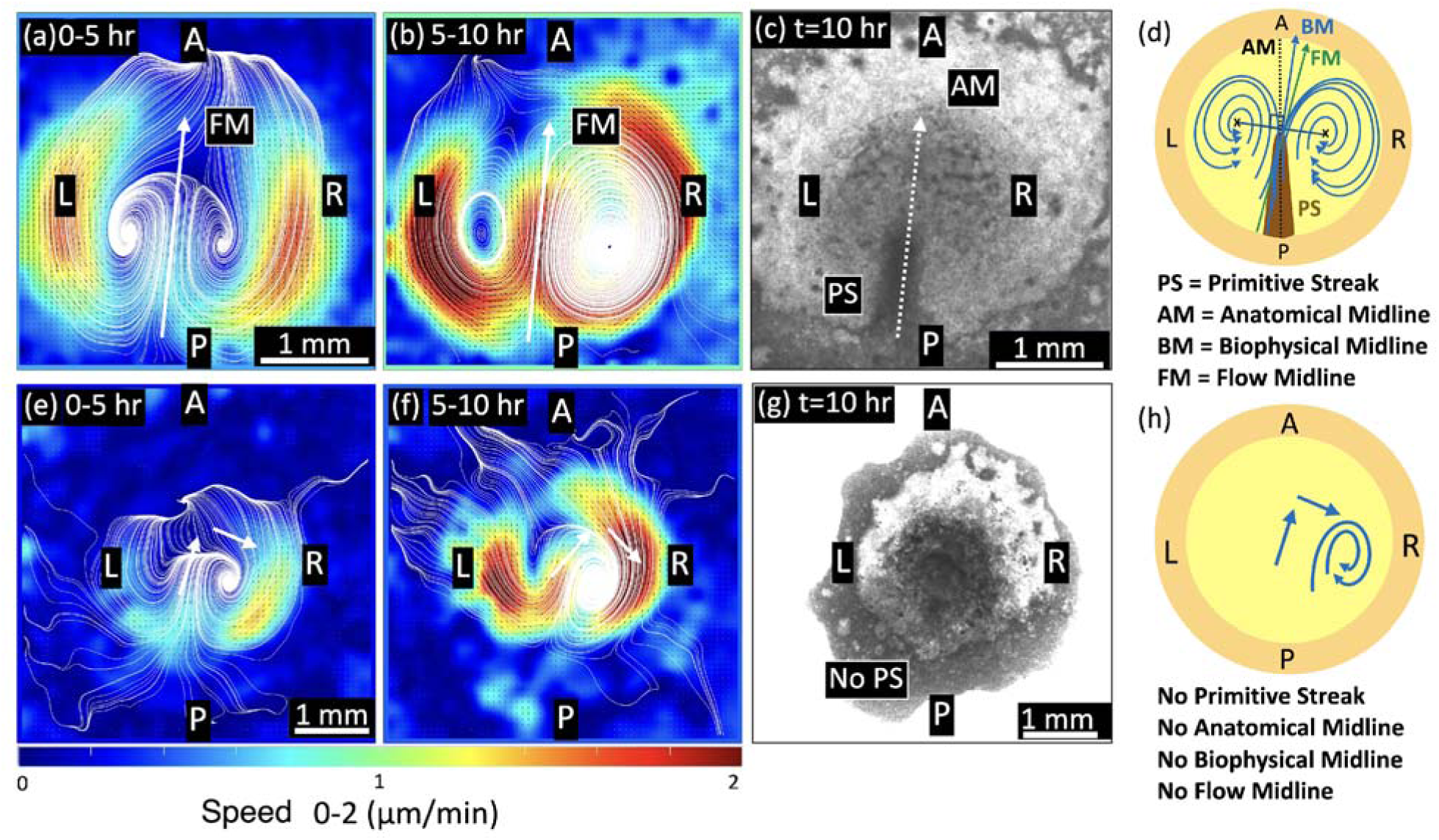
Comparison of cellular flows in early chicken embryo development: control vs cell division inhibited. Time-averaged (5 hour) speed results from PIV analysis of the control and aphidicolin-treated embryos (see Table 1). **Top panels – control embryo** (control sample 1, Table 1): (a) During the initial period (0-5 hours), the bilateral flows have a high degree of L-R symmetry. (b) In the later period (5-10 hours), the bilateral flows may still be maintained but the L-R symmetry is lost, and the R dominance is visible. (c) The brightfield image at the end time point (t = 10 hours) showing PS. (d) The schematic represents the bilateral cellular flows in the control, with flows dominant in R. The PS forms but the AM and BM axes may have a slight deviation from each other. **Bottom panels – cell-division inhibited embryos** (aphidicolin sample 3, Table 1): (e) During the initial period (0-5 hours), there is a single vortex formation on the R side. (f) In the later period (5-10 hours), the single R vortex persists and dominates the flow speeds. (g) The brightfield image at the end time point (t = 10 hours) with no PS formation. (h) The schematic represents the cellular flows in cell-division inhibited embryos, with single R vortex dominated flows. The PS did not form, and the midlines AM, BM, and FM were not defined. The short arrows serve as a guide to indicate direction of the flows at that location, not representing the midlines (Methods). The length scale bar in (a) is the same for all panels in top row, and length scale bar in (e) is same for all panels in bottom row.

Looking at the half-time averaged cellular flow speeds in the control case, in the first half (0-5 hours), the flows were symmetric in L and R (Figure 6a). Whereas, in the second half (5-10 hours), the flows became R dominant (Figure 6b), demonstrating that bilateral flows end with a R dominance during early midline morphogenesis (Figure 6a, b). In these embryos, the bilateral cellular flows occurred during the formation of the PS structure (Figure 6c), and the three distinct midlines (AM, BM, and FM) can be defined, tracked, and quantified.

In the cell division inhibited case, the cellular flows deviated from the classic bilateral mode (Figure 6 e, f). R dominance of the cellular flows was recognizable in both the first half (0-5 hours, Figure 6e) and the second half (5-10 hours, Figure 6f), with a single R vortex forming towards the end. The three distinct midlines were not defined because of these complex cellular flow patterns and diminished PS formation (Figure 6g) under mitotic arrest. Taken together, R dominance of the bilateral cellular flows persisted even though PS formation was diminished under mitotic arrest.

## Discussion

The present study identified previously undetected LR asymmetries in the bilateral cellular flows (the polonaise movements) during early midline morphogenesis in chick gastrulation. Although the bilateral cellular flows have been studied for approximately a hundred years (10, 11, 30-35), this is the first time LR asymmetries in flow have been identified. Current models assume that the LR laterality in amniotes arises in later stages of development, when the set of the well-known laterality genes triggers asymmetry patterning after the appearance of the node (4, 6, 27, 36-38). An implication, based on the current model, would be that earlier patterning events, including the bilateral cellular flows, are symmetric. However, our results are inconsistent with this assumption. The present study with detailed biophysical quantification techniques and novel approaches for time-averaging of cell speeds and vorticity reveals that bilateral flows had already exhibited LR asymmetry towards R dominant flow during PS extension and prior to Hensen’s node formation. Thus, our unique time-averaging approach to quantify bilateral cellular flows enabled the visualization and quantification of these LR flow asymmetries.

Our present study showed that the LR asymmetric expression of the laterality genes became remarkable at Hensen’s node by WISH with NBT/BCIP staining (Figure 3, Figure S3). This classical method has been generally utilized in previous studies to examine the expression of the laterality genes (4, 6, 27, 36-38), however, its sensitivity of RNA detection is relatively lower than the latest techniques such as HCR (28) and Spatial transcriptomics (39). These higher sensitivity/resolution techniques could help to examine the spatio-temporal gene expression pattern in higher resolution and to reveal unknown mechanism(s) of gene regulation in the cellular flows.

Our experimental approaches demonstrated that mitotic arrest maintained the cellular flows despite diminished PS formation and showed initial bilateral movements that later transitioned to R dominance and a single vortex. These results suggest that PS formation is not crucial for initiating the LR laterality in the cellular flows. Our data support the model in which the LR asymmetry in amniote gastrula is already set up, as detected by the cellular flow patterns, prior to when the node-dependent LR patterning kicks in.

If the bilateral cellular flows were symmetric, the three distinct midline axes (AM, BM, and FM, shown in Figure 4 and Figure S4) were expected to overlap completely. At the onset of the cellular flow, there was a significant difference in positions for the three midlines. The reason for the high variation is currently unknown but perhaps due to the flow not being fully developed in the beginning (first few hours). The angular separation between the midlines starts reducing as the flow becomes stable as PS continues to extend, but a small difference persists. These temporal variations in the flow midlines could be an important factor that gives rise to the initial LR asymmetry in the bilateral flows. Interestingly, the AM and PS maintain a linear structure during extension despite the LR imbalanced cellular flow (Figure1, Figure 3, Movie1, and Figure S3) (22, 40). Our previous study has shown that the authentic PS extends regardless of aberrant flow patterns caused by ectopically induced Vg1 (an axis-inducing morphogen) and secondary PS formation (29). These data support each other and imply that the bilateral rotational cellular flow is not necessarily responsible for PS extension. Rather, the PS is capable of extending by internal mechanism(s), such as convergent extension within the PS (18, 33, 34). Our results imply that the emergent flow mechanics that arise from collective cell movements break LR symmetry and impart LR laterality much earlier than previously expected.

Another unpredicted finding of our study is the R dominance of the flows at later stages during PS extension. The signature of the R dominance was seen readily in the control embryos (Figure 2 and 4, Figure S7-S10). In the mitotic arrest case, the cellular flows are highly asymmetric (non-bilateral). The absence of the PS structure may remove or reduce mechanical stability. Consistent with this idea, in some cases, the flows started to develop bilateral-type movements but merged on one side. In all cases with mitotic arrest, this merging flow was always towards R by forming a single vortex in our data (N=4). This implies that the R dominance already present in control embryos is being further amplified in the mitotic arrest cases. It remains to be explored how the preferred R dominance is induced and or patterned.

Two models have been proposed for establishing the LR laterality in the chick embryo (8, 41). The classic model is that the LR laterality is set up in the blastoderm by localization of maternal determinant before gastrulation and each LR side develops by following the pre-set pattern(s) (41). The recent model is that molecular signaling(s) from the midline structures, including the PS, initiates and patterns the LR laterality (8) based on extensive genetic and molecular studies (6, 27, 42). Our present study with quantitative biophysical analyses has revealed that the LR asymmetry in the bilateral cellular flow appears earlier than remarkable asymmetries in the expression of the LR regulatory genes at the Hensen’s node (Figure 3, Figure S3). Further, the R dominance in the flow pattern was preserved even though the PS, at the anterior end of which Hensen’s node arises, was diminished (Figure 5). These data suggest that the LR laterality in the cellular flow is unlikely to depend on the node-driven LR patterning signaling(s). It is currently unknown how the cellular flow is induced and how the LR asymmetry in the bilateral cellular flow is patterned. However, the chirality of movement in mammalian cells has been extensively studied in vitro and appears to be intrinsic and readily scaled up to the level of tissues in bioengineered multicellular systems (43-52). Relevant phenomena with the chirality of cytoskeletons have been identified in invertebrates at early developmental stage prior to LR-organizer formation and play important roles in LR-body patterning (53-56). These previous studies imply that the cellular chirality, exhibited by their movement and cytoskeleton, may be evolutionarily involved in LR-body patterning (57, 58). In the chick embryos at the pre-streak stage, several morphogens (e.g. Wnts and Vg1) are already expressed in the embryonic disc prior to and at the onset of cellular flows. Whereas these morphogens are known as axis-inducing factors (59), their roles on the large-scale cellular flow, particularly in amniotes, remain to be determined. We have previously demonstrated the ectopic introduction of Vg1 modulates the cellular flow pattern (29). These morphogens would regulate the cellular chirality, generating the LR asymmetry in the bilateral cellular flow. Further studies examining any LR asymmetries of these morphogens and/or the response in single cells may reveal undefined mechanism(s) of the LR laterality of the cellular flow, such as the early symmetry breaking.

While large-scale cellular flows during gastrulation are evolutionarily conserved (60-62), the LR laterality of the cellular flows remains largely unexplored in most species. Chick embryos begin the cellular flow with the bilateral rotating pattern prior to and during early PS extension and then shift to a lateral movement toward the PS as the ingression takes place at the midline for germ layer formation (31, 32). We have previously reported that the midline structures, including the PS, are required for both LR asymmetry and the ipsilaterality of the lateral cell movements, which subsequently occur after HH3 following the polonaise movements (19). The current study has demonstrated that mitotic arrest diminished PS formation but the LR laterality in the bilateral rotating cellular flow can start (Figure 5 and Figure S12). These findings suggest that the role of the midline structures for the laterality/ipsilaterality of the cellular flows would switch in cellular flow patterning the early and later developmental windows of the large-scale cellular flow.

Whether the LR laterality of the cellular flow is related to the later genetically programmed LR patterning remains to be determined. The asymmetric expression of the LR regulatory genes initiates at the Hensen’s node (also termed as the ‘node’ in mice, Figure 3) (25-27). In contrast, the Hensen’s node in chick gastrula has short primary cilia instead the motile cilia (63, 64). These insights suggest that the model of the cilia-regulating gene expression may not be applicable in all amniotes, and alternative models have been proposed (20, 63, 65). Further studies, for example, disruption of the asymmetric counter-rotating cellular flow, will clarify the role of the cellular flows for the genetically programmed LR patterning. Our study would provide an alternative view for symmetry breaking and provides hints for understanding the establishment of LR laterality and bilateral body patterning.

## Materials and Methods

### Embryo isolation and culture condition

We obtained fertilized eggs (White Leghorn; *Gallus Gallus domesticus*) from Petaluma farms in California. From unincubated eggs, pre-streak stage embryos [stage X to XII (24)] were isolated in Tyrode’s solution (final concentration: 137mM NaCl, 2.7 mM KCL, 1mM MgCl2, 1.8mM CaCl2, 0.2 mM Ha2HPO4, 5.5mM D-glucose, pH 7.4). The chick embryos were subsequently cultured until HH3 for about 14 hours at 37°C, by using in vitro culture method, termed as ‘New culture’ (66) as previously described (29, 40, 66, 67). After electroporation, the embryos with reagents (described as below) were live-imaged on a vitelline membrane stretched around a glass ring according to the New culture method as previously described (29).

### Electroporation

Isolated embryos from the yolk were transfected with control-oligo DNA (5’-CCTCTTACCTCAGTTACAATTTATA-3) conjugated with Lissamine (GENE TOOLS) using an electroporator (Nepagene) with 3 pulses of 2.4-3.8 Volts, 50 milliseconds duration, 500 milliseconds interval, and platinum electrodes. The DNA solution delivered to the epiblast contained 0.1% fast green (final 0.02%), 80% glucose (final 4%) and 1mM control-oligo.

### Aphidicolin treatment

Aphidicolin (SIGMA, A0781), dissolved in DMSO (SIGMA, B23151), was added to Tyrode’s solution to a final concentration of 100μM. The isolated embryos were soaked in Tyrode’s solution with either 0.3% DMSO (control), or 100μM aphidicolin for 15 minutes at 37°C. Embryos were then cultured using the New culture method at 37°C.

### BrdU assay

Embryos were soaked in Tyrode’s solution containing BrdU (final concentration; 0.1mM, Thermo Fisher, B23151) with or without aphidicolin (SIGMA, A0781) at 37°C for 15 minutes, and cultured for 16 hours at 37°C (BrdU was incorporated for 16 hours). Embryos were then fixed in 4% paraformaldehyde (PFA, Electron Microscopy Sciences)/PBS for 30 minutes at room temperature (RT). Embryos were then washed with PBS to remove PFA, and unincorporated BrdU, and incubated with 1M HCl for 1 hour at RT to denature DNA. BrdU signal was detected by immunofluorescence staining with anti-BrdU antibody (1∶200, Millipore, MAB3424).

### Whole-mount In Situ Hybridization (WISH)

The detailed ISH protocol was described in previous studies (19, 29). The cultured embryos were fixed in 4% PFA at 4°C, and washed in PBST for 5 minutes in 3 times. After a sequential methanol replacement from 25% to 100% and a subsequent rehydration to PBST, the embryos were treated with proteinase K solution (final concentration: 0.5mg/mL)/PBST for 1min at RT. The reaction was neutralized by glycine (2mg/mL)/PBST and washed in PBST for 5 minutes in two times. After 2 hours of pre-hybridization in Solution I (final concentration: 1% SDS, 5xSSC pH4.5, 50% formamide) with yeast tRNA (final concentration: 50μg/mL, MilliporeSigma, MA) and heparin (final concentration: 50μg/mL, MilliporeSigma, MA), the embryos were hybridized with RNA-probes at 70°C overnight [probes; *Brachyury, Sox3* (a gift from Dr. Raymond B. Runyan), *Slug, Lefty, Nodal, Shh*]. Next day, the buffer was replaced to new Solution I, and the embryos were washed for 30 minutes two times. Next, washed in Solution III (final concentration: 50% formamide, 2XSSC pH4.5) for 30 minutes two times, followed by washing in TBST at RT, 3 times for 5 minutes each and blocking with 10% sheep serum in TBST at RT. Anti-DIG antibody (final concentration: 1/2000, Roche, MA) was reacted with the embryos at 4°C overnight. Then, the following day, embryos were washed in TBST for 1 hour, 5 times at RT. After treatment with NTMT (final concentration: 100mM of NaCl, 100mM of 50mM MgCl2, 0.1% Triton) twice for 5 minutes, color-development was performed with NBT/BCIP solution in NTMT (4.5μL of NBT and 3.5μL of BCIP in 1mL of NTMT). All ISH experiments were parallelly carried out with control embryos to manage the appropriate time of color-development.

### Imaging of the samples

Live-imaging of the fluorescently-labeled chick embryos was performed at 37C° on either microscopes, the Widefield Epi-fluorescent inverted microscope (Nikon Ti inverted fluorescent Microscope with CSU-W1 large field of view, UCSF Nikon imaging center) connected by ANDOR iXon camera or the TIRF/Spinning Disk microscope (Nikon Ti inverted fluorescent Microscope with CSU-22 spinning disc confocal, UCSF Nikon imaging center) connected with Prime 95B Scientific CMOS camera. After live-imaging (29), these samples were subsequently subjected to ISH and imaged by the Leica MZ16F with DFC300 Fx camera and FireCam V.3.4.1 software. BrdU-stained samples were imaged on Ti2 inverted fluorescence microscope connected by the Crest LFOV spinning disk confocal at UCSF Nikon imaging center.

### Image Processing

Time-lapse imaging datasets from the microscope were opened in the ImageJ (or Fiji) software and split into two channels: brightfield and fluorescence. The brightness and contrast settings of the fluorescence time-lapse images were optimized using suitable thresholds to reduce the noise and enhance the ability to clearly resolve the bright fluorescent markers against a dark background. In the time-lapse sequence, the cellular movements in the embryo only start after a short time delay. For our biophysical motility analysis, we only consider those images after the start of cell movements, this starting point would correspond to the zero time (t), or t = 0 in our analysis. In all the datasets, after determining the starting time, we analyze time-lapse images corresponding to the first 10 hours of early development. This helps make standard time-point comparisons between the different experimental datasets presented here.

### Particle Image Velocimetry (PIV)

PIV is an experimental technique to quantify fluid flow-fields. This non-intrusive technique is traditionally based on optical time-lapse imaging of tracer particles that are typically illuminated by a high-intensity light source. Next, the time-lapse image sequence is analyzed further and processed using computational algorithms to obtain a spatiotemporal map of flow velocity vectors. Here, we have utilized PIV analysis techniques on the time-lapse images of fluorescently tagged cells in the embryo to quantify their movements and speeds over time. We employed the MATLAB-based PIVlab package for our data analysis (21) (Supporting Information).

### Time averaging of PIV datasets

The PIV analysis by default provides instantaneous velocity vectors that are calculated between two time-points (here this timescale is 3 min). In our datasets, the cell movements over short timescales of 3 mins are very small and the time-resolved transitions in the datasets are inherently noisy due to spatial and temporal fluctuations. Hence, we calculated the time-averaged measurements of the key cell movement parameters (speeds, vorticity) over longer timescales. We found that these time-averaged measurements were significantly better for both detailed quantification and interpretation of asymmetry in the bilateral cellular flows. We carried out three different durations of time averaging – (i) 10 hours, (ii) 5 hours, and (iii) 1 hour. This enables interpretation and comparison of bilateral cellular flows at different timescales in the datasets. (i) Time averaging over 10 hours gives an overall picture of the cellular flows during early development. (ii) Time averaged quantification over 5 hours enables us to distinguish between the first half and second half of the process. (iii) Time averaged quantification over 1 hour enables us to carefully study questions about cellular flows such as spatial and temporal origins, changes and transitions, and dominance of L-R asymmetry spanning 10 time-points distributed over every 1 hour. The two key parameters that we use to quantify and characterize cellular flows are speed and vorticity, and we have studied their variation over time during the process of development.

### Speed plots

The speed at every spatial point at a given time is calculated as the vector magnitude of the x and y components of the velocities in PIVlab. The cellular flow speeds are plotted based on the thresholds (described below) at each time point for all the datasets using a heatmap color scheme continuously varying from low speeds (blue) to higher speeds (red).

### Streamline plots

Streamlines are the instantaneous and simplified representations of fluid flow directions or paths. Each streamline is tangent to the velocity vector of the fluid flow at every point along its path. They are very helpful in visualizing and analyzing fluid flow. The flow parameters such as vorticity can be better interpreted by comparison with the streamlines. Here, the streamlines were plotted manually with appropriate line width and color. Selecting the “draw streamlines” option in PIVlab and clicking on the vectors automatically generates the most useful streamlines.

### Vorticity plots

Vorticity is a vector quantity commonly used in fluid dynamics to quantify the local rotation of a fluid element. Mathematically, the vorticity is calculated as the curl of the fluid velocity. It provides important insights into the rotational regions in the cellular flows, and this rotation can either be clockwise or counterclockwise. High vorticity regions indicate strong local rotation, whereas low vorticity points towards weak rotation. Vorticity plots were generated for all datasets in PIVlab using a heatmap color scheme continuously varying from high clockwise vorticity (red) to 0 (white) to high counterclockwise vorticity (blue).

### Biophysical quantification of PIV datasets

In addition to the default plotting of PIV velocity flow fields and their derived quantities such as speed and vorticity, we further extracted quantitative information to characterize the cellular flows in different datasets as described below (Supporting Information).

### Midlines

We defined three midlines in the control datasets: AM (defined by the biological structure), BM and FM (defined by the cellular flows). The Anatomical Midline (AM) is the line passing through the center of Primitive streak (PS) (Figure 1a). The bilateral flow establishes two counter-rotating vortices in L and R regions. The line drawn perpendicular to the midpoint of the line joining the vortex centers is defined as Biophysical Midline (BM) (Figure S4a). A line drawn parallel to the flow streamlines passing through the midpoint of the vortex centers is defined as the Flow Midline (FM) (Figure S4a) (Supporting Information).

### Vorticity and speed thresholds for quantification

In each control sample dataset, for every hourly averaged speed plot, the maximum speed (V_max_) value was identified. Based on this V_max_ value, three thresholds of 30%, 50% and 70 % were calculated (Table S3). Here, 30% threshold means velocity vectors with the magnitude of 30% of V_max_ (i.e. V_max_*0.3) were retained, and similarly for 50% and 70%. Similarly for vorticity, the maximum vorticity (magnitude) value was obtained over one hour. Again, based on this maximum vorticity magnitude the three thresholded values, 30% 50% and 70% were obtained (Table S4). Speed and vorticity values that are higher than these thresholds were considered for quantifying the areas on the left (A_L_) and right (A_R_) regions. This approach enabled a rigorous quantification of cellular flows at different magnitudes (with varying thresholds in speed and vorticity) and the comparison between L and R regions over time (see Figure S7-S10, supporting information).

### Statistical Test

Statistical analysis was done using the non-parametric Wilcoxon test in JMP Pro 17.0.0. The obtained p-values are mentioned in the relevant figure legends and in the supplementary tables (Table S5 and S6).

## Acknowledgments

We thank present and past Mikawa lab and Prakash lab members for their invaluable suggestions and/or assistance. We also thank Monu Verma, Santhan Chandragiri, and Melissa Ruszczyk for helpful discussions and suggestions. This work was supported in part by grants from NIH (R01HL122375, R37HL078921, R01HL132832, R01HL148125) to T.M.; Uehara Memorial Foundation Fellowship and JSPS Postdoctoral Fellowship for Research Abroad to R.A.; University of Miami for startup funding support to V.N.P.

## References

1. M. Q. Martindale, J. R. Finnerty, J. Q. Henry, The Radiata and the evolutionary origins of the bilaterian body plan. Molecular Phylogenetics and Evolution 24, 358–365 (2002).

2. E. K. Namigai, N. J. Kenny, S. M. Shimeld, Right across the tree of life: the evolution of left-right asymmetry in the Bilateria. Genesis 52, 458–470 (2014).

3. J. P. Richter, R. C. Bell, The Notebooks of Leonardo Da Vinci (Dover Publications, 1970).

4. S. Nonaka et al., Randomization of left-right asymmetry due to loss of nodal cilia generating leftward flow of extraembryonic fluid in mice lacking KIF3B motor protein. Cell 95, 829–837 (1998).

5. J. J. Essner, J. D. Amack, M. K. Nyholm, E. B. Harris, H. J. Yost, Kupffer’s vesicle is a ciliated organ of asymmetry in the zebrafish embryo that initiates left-right development of the brain, heart and gut. Development 132, 1247–1260 (2005).

6. C. J. Tabin, The key to left-right asymmetry. Cell 127, 27–32 (2006).

7. M. Levin, N. Nascone, Two molecular models of initial left-right asymmetry generation. Med Hypotheses 49, 429–435 (1997).

8. N. A. Brown, L. Wolpert, The development of handedness in left/right asymmetry. Development 109, 1–9 (1990).

9. M. Chuai et al., Cell movement during chick primitive streak formation. Dev Biol 296, 137–149 (2006).

10. M. Saadaoui, D. Rocancourt, J. Roussel, F. Corson, J. Gros, A tensile ring drives tissue flows to shape the gastrulating amniote embryo. Science 367, 453–458 (2020).

11. J. Firmino, D. Rocancourt, M. Saadaoui, C. Moreau, J. Gros, Cell Division Drives Epithelial Cell Rearrangements during Gastrulation in Chick. Dev Cell 36, 249–261 (2016).

12. G. Serrano Nájera, C. J. Weijer, Cellular processes driving gastrulation in the avian embryo. Mech Dev 163, 103624 (2020).

13. L. Bodenstein, C. D. Stern, Formation of the chick primitive streak as studied in computer simulations. J Theor Biol 233, 253–269 (2005).

14. S. A. Sandersius, M. Chuai, C. J. Weijer, T. J. Newman, A ‘chemotactic dipole’ mechanism for large-scale vortex motion during primitive streak formation in the chick embryo. Phys Biol 8, 045008 (2011).

15. B. Vasiev, A. Balter, M. Chaplain, J. A. Glazier, C. J. Weijer, Modeling gastrulation in the chick embryo: formation of the primitive streak. PLoS One 5, e10571 (2010).

16. M. Serra et al., A mechanochemical model recapitulates distinct vertebrate gastrulation modes. Sci Adv 9, eadh8152 (2023).

17. M. Chuai, G. Serrano Nájera, M. Serra, L. Mahadevan, C. J. Weijer, Reconstruction of distinct vertebrate gastrulation modes via modulation of key cell behaviors in the chick embryo. Sci Adv 9, eabn5429 (2023).

18. O. Voiculescu, L. Bodenstein, I. J. Lau, C. D. Stern, Local cell interactions and self-amplifying individual cell ingression drive amniote gastrulation. Elife 3, e01817 (2014).

19. L. Maya-Ramos, T. Mikawa, Programmed cell death along the midline axis patterns ipsilaterality in gastrulation. Science 367, 197–200 (2020).

20. D. S. Adams et al., Early, H+-V-ATPase-dependent proton flux is necessary for consistent left-right patterning of non-mammalian vertebrates. Development 133, 1657–1671 (2006).

21. W. Thielicke, E. J. Stamhuis, PIVlab – Towards User-friendly, Affordable and Accurate Digital Particle Image Velocimetry in MATLAB. Journal of Open Research Software 2 (2014).

22. V. Hamburger, H. L. Hamilton, A series of normal stages in the development of the chick embryo. 1951. Dev Dyn 195, 231-272 (1992).

23. W. Thielicke, R. Sonntag, Particle Image Velocimetry for MATLAB: Accuracy and enhanced algorithms in PIVlab. (2021).

24. H. Eyal-Giladi, S. Kochav, From cleavage to primitive streak formation: a complementary normal table and a new look at the first stages of the development of the chick. I. General morphology. Dev Biol 49, 321–337 (1976).

25. K. Katsu, D. Tokumori, N. Tatsumi, A. Suzuki, Y. Yokouchi, BMP inhibition by DAN in Hensen’s node is a critical step for the establishment of left-right asymmetry in the chick embryo. Dev Biol 363, 15–26 (2012).

26. T. Mikawa, A. M. Poh, K. A. Kelly, Y. Ishii, D. E. Reese, Induction and patterning of the primitive streak, an organizing center of gastrulation in the amniote. Dev Dyn 229, 422–432 (2004).

27. M. Levin, R. L. Johnson, C. D. Stern, M. Kuehn, C. Tabin, A molecular pathway determining left-right asymmetry in chick embryogenesis. Cell 82, 803–814 (1995).

28. H. M. Choi et al., Programmable in situ amplification for multiplexed imaging of mRNA expression. Nat Biotechnol 28, 1208–1212 (2010).

29. R. Asai, V. N. Prakash, S. Sinha, M. Prakash, T. Mikawa, Coupling and uncoupling of midline morphogenesis and cell flow in amniote gastrulation. eLife 12, RP89948 (2024).

30. C. Cui, X. Yang, M. Chuai, J. A. Glazier, C. J. Weijer, Analysis of tissue flow patterns during primitive streak formation in the chick embryo. Dev Biol 284, 37–47 (2005).

31. R. Wetzel, Untersuchungen am Hühnchen. Die Entwicklung des Keims während der ersten beiden Bruttage. Wilhelm Roux Arch Entwickl Mech Org 119, 188–321 (1929).

32. L. Gräper, Die Primitiventwicklung des Hühnchens nach stereokinematographischen Untersuchungen, kontrolliert durch vitale Farbmarkierung und verglichen mit der Entwicklung anderer Wirbeltiere. Wilhelm Roux Arch Entwickl Mech Org 116, 382–429 (1929).

33. O. Voiculescu, F. Bertocchini, L. Wolpert, R. E. Keller, C. D. Stern, The amniote primitive streak is defined by epithelial cell intercalation before gastrulation. Nature 449, 1049–1052 (2007).

34. E. Rozbicki et al., Myosin-II-mediated cell shape changes and cell intercalation contribute to primitive streak formation. Nat Cell Biol 17, 397–408 (2015).

35. P. Caldarelli et al., Self-organized tissue mechanics underlie embryonic regulation. Nature 633, 887–894 (2024).

36. T. Nakamura, H. Hamada, Left-right patterning: conserved and divergent mechanisms. Development 139, 3257–3262 (2012).

37. Y. Okada, S. Takeda, Y. Tanaka, J. I. Belmonte, N. Hirokawa, Mechanism of nodal flow: a conserved symmetry breaking event in left-right axis determination. Cell 121, 633–644 (2005).

38. K. Minegishi et al., A Wnt5 Activity Asymmetry and Intercellular Signaling via PCP Proteins Polarize Node Cells for Left-Right Symmetry Breaking. Dev Cell 40, 439–452 e434 (2017).

39. P. L. Ståhl et al., Visualization and analysis of gene expression in tissue sections by spatial transcriptomics. Science 353, 78–82 (2016).

40. R. Bellairs, M. Osmond, Atlas of chick development (Elsevier, 2005).

41. H. Wilhelmi, Experimentelle Untersuchungen über Situs inversus viscerum. Archiv für Entwicklungsmechanik der Organismen 48, 517–532 (1921).

42. S. M. Pagán-Westphal, C. J. Tabin, The transfer of left-right positional information during chick embryogenesis. Cell 93, 25–35 (1998).

43. X. Yao, X. Wang, J. Ding, Exploration of possible cell chirality using material techniques of surface patterning. Acta Biomater 126, 92–108 (2021).

44. Y. Bao, et al., Early Committed Clockwise Cell Chirality Upregulates Adipogenic Differentiation of Mesenchymal Stem Cells. Adv Biosyst 4, e2000161 (2020).

45. S. Utsunomiya et al., Cells with Broken Left–Right Symmetry: Roles of Intrinsic Cell Chirality in Left–Right Asymmetric Epithelial Morphogenesis. Symmetry 11, 505 (2019).

46. H. K. Kwong, Y. Huang, Y. Bao, M. Lam, T. H. Chen (2018) Cell Chiral Orientation Enhanced by Intercellular Alignment. in 2018 IEEE 12th International Conference on Nano/Molecular Medicine and Engineering (NANOMED), pp 104-108.

47. J. Fan et al., Cell chirality regulates intercellular junctions and endothelial permeability. Sci Adv 4, eaat2111 (2018).

48. L. Q. Wan, A. S. Chin, K. E. Worley, P. Ray, Cell chirality: emergence of asymmetry from cell culture. Philos Trans R Soc Lond B Biol Sci 371 (2016).

49. A. Dimonte, A. Adamatzky, V. Erokhin, M. Levin, On chirality of slime mould. Biosystems 140, 23–27 (2016).

50. H. Yamanaka, S. Kondo, Rotating pigment cells exhibit an intrinsic chirality. Genes Cells 20, 29–35 (2015).

51. T. Yamamoto et al., Epithelial cell chirality emerges through the dynamic concentric pattern of actomyosin cytoskeleton. bioRxiv 10.1101/2023.08.16.553476, 2023.2008.2016.553476 (2024).

52. A. Tamada, S. Kawase, F. Murakami, H. Kamiguchi, Autonomous right-screw rotation of growth cone filopodia drives neurite turning. J Cell Biol 188, 429–441 (2010).

53. V. N. Meshcheryakov, L. V. Beloussov, Asymmetrical rotations of blastomeres in early cleavage of gastropoda. Wilehm Roux Arch Dev Biol 177, 193–203 (1975).

54. M. Inaki, T. Sasamura, K. Matsuno, Cell Chirality Drives Left-Right Asymmetric Morphogenesis. Front Cell Dev Biol 6, 34 (2018).

55. M. Inaki, J. Liu, K. Matsuno, Cell chirality: its origin and roles in left-right asymmetric development. Philos Trans R Soc Lond B Biol Sci 371 (2016).

56. M. Inaki et al., Chiral cell sliding drives left-right asymmetric organ twisting. Elife 7 (2018).

57. L. N. Vandenberg, M. Levin, Far from solved: a perspective on what we know about early mechanisms of left-right asymmetry. Dev Dyn 239, 3131–3146 (2010).

58. L. N. Vandenberg, M. Levin, A unified model for left-right asymmetry? Comparison and synthesis of molecular models of embryonic laterality. Dev Biol 379, 1–15 (2013).

59. S. B. Shah et al., Misexpression of chick Vg1 in the marginal zone induces primitive streak formation. Development 124, 5127–5138 (1997).

60. S. F. Gilbert, M. J. F. Barresi, Developmental Biology (Oxford University Press, 2017).

61. L. Solnica-Krezel, D. S. Sepich, Gastrulation: making and shaping germ layers. Annu Rev Cell Dev Biol 28, 687–717 (2012).

62. M. Leptin, Gastrulation movements: the logic and the nuts and bolts. Dev Cell 8, 305–320 (2005).

63. L. A. Stephen et al., The chicken left right organizer has nonmotile cilia which are lost in a stage-dependent manner in the talpid(3) ciliopathy. Genesis 52, 600–613 (2014).

64. J. Männer, Does an equivalent of the “ventral node” exist in chick embryos? A scanning electron microscopic study. Anat Embryol (Berl) 203, 481–490 (2001).

65. J. Gros, K. Feistel, C. Viebahn, M. Blum, C. J. Tabin, Cell movements at Hensen’s node establish left/right asymmetric gene expression in the chick. Science 324, 941–944 (2009).

66. D. A. T. New, A New Technique for the Cultivation of the Chick Embryo in vitro. Journal of Embryology and Experimental Morphology 3, 326–331 (1955).

67. R. Asai, M. Bressan, T. Mikawa, Avians as a Model System of Vascular Development. Methods Mol Biol 2206, 103–127 (2021).

