## Supporting Information for "Bilateral cellular flows display asymmetry prior to left-right organizer formation in amniote gastrulation"

[1]Cardiovascular Research Institute, University of California, San Francisco. San Francisco, California 94158, USA. [2]Department of Physics, University of Miami, Coral Gables, Florida 33146, USA. [3]Department of Biology, University of Miami, Coral Gables, Florida 33146, USA. [4]Department of Marine Biology and Ecology, University of Miami, Miami, Florida 33149, USA. [5]Kumamoto University, IRCMS, Kumamoto, 860-0811, Japan.

**Equal contribution

*Corresponding authors: Vivek N. Prakash, Takashi Mikawa

**This PDF file includes:**

Supporting text
Figures S1 to S12
Tables S1 to S6
Legends for Movies S1 to S2

**Other supporting materials for this manuscript include the following:**

Movies S1 to S2

### Methods

**Particle Image Velocimetry.** The first step of the PIV analysis is to input the brightness optimized fluorescence image/frame sequence into the PIVlab package choosing the time-resolved sequence option. The images were pre-processed using the following settings: high pass kernel size - 15 pixels, and wiener2 denoise filter window size – 3 pixels. Next, we proceed to do the PIV cross-correlation analysis on the preprocessed images using the FFT window deformation algorithm. The interrogation window sizes were chosen to be 128 x 128 pixels with 50% overlap for pass 1 and 64 x 64 pixels with 50% overlap for pass 2. Most of the datasets were analyzed using the above-mentioned settings, but there were variations for a few datasets (see Table S1). After completing this analysis step, we obtain the raw time-resolved velocity vector fields. Next, we carry out post processing of these velocity vector fields to reduce the noise by selecting suitable velocity limits, standard deviation, and local median filters. After this post-processing step, the time-resolved velocity vector fields, the velocity magnitudes (speeds), and vorticity parameters were extracted for each image/frame.

**Midlines.** A line drawn parallel to the flow streamlines passing through the midpoint of the vortex centers is defined as the Flow Midline (FM) (Fig. S4a). In the PIVlab GUI, streamlines were drawn for each hourly averaged vector frame. The images with streamlines over the vectors were imported in ImageJ. Next, straight lines were drawn approximately parallel to the streamlines passing near the midpoints of the vortex centers. The average of these straight lines was considered as the FM. The coordinates lying at the two opposite ends of the FM in the image were noted. Then the PIV MATLAB file containing hourly averaged vector information for a sample was loaded into memory. The recorded coordinates on FM were used to draw a straight line. Next, the grids which lay on or near the straight line were set to zero, dividing the PIV matrix into two regions, L and R about the FM. This process was repeated in all the 10 hourly averaged frames for one sample.

**Temporal analysis.** For speed area quantification in control embryos, in each sample, we obtained the maximum velocity magnitude, $V_{max}$, over an hourly time window (Table S3). Based on the value of $V_{max}$, three threshold values were chosen: 30%, 50% and 70%. Here, a 30% threshold represents the velocity magnitude of $0.3*V_{max}$, similarly for 50% and 70% (Table S3). For every time averaged window, velocity vector fields with magnitudes greater than or equal to these thresholds (for example 50% corresponds to $0.5*V_{max}$) were retained in the L and R regions about the midline. Next, we calculated the number of vectors which were above the cut-off values ($0.5*V_{max}$ for 50%) in the L and R regions. Since each vector occupies one grid, these number of vectors were rescaled to represent the actual area in the L and R regions. This process is repeated for every time-averaged plot with its own $V_{max}$ for every sample (Table S3).

For vorticity area quantification, the same quantification is performed by obtaining the maximum vorticity magnitude in each hourly averaged window and comparing L and R sides based on the three thresholds (30%, 50% and 70%) of the maximum value (see Table S4).

The three thresholds (30%, 50% and 70%) (low, medium, high) helped in quantitatively comparing cellular flows in L and R regions at different flow strength levels (speeds). For example, a threshold of 30% in speed allowed more velocity vectors (vectors with a magnitude higher than $0.3*V_{max}$) to contribute to the speed area in L and R. In the 30% case, we observed that the L-R speed areas were symmetric (see Figure S10), i.e. there was no significant differences between the L and R sides. Whereas, with a stricter threshold of 70% (vectors with a magnitude higher than $0.7*V_{max}$) led to the selection of a lesser number of velocity vectors (hence, lesser area) which showed an asymmetric distribution of L and R areas (R dominated, see Figure S10). We chose to display results of the speed and vorticity areas using the medium threshold (50%) in Figure 4. For the low and high thresholds (30% and 70%), we do not expect the results to change significantly if we had chosen different values (say 25% and 75%).

**Supplementary Movies**

**Movie1**: **Cellular flows reveal L-R asymmetry during early chick embryo development.**
The top left panel is a brightfield imaging time-lapse video of early chick embryo development, revealing the development of the Primitive Streak (PS). The top right is a Flowtrace visualization video showing large-scale bilateral cellular flows or 'polonaise movements', the cells are tagged with fluorescent labels. The bottom left is a heatmap video of cellular speeds, calculated using the Particle Image Velocimetry (PIV) technique. The bottom right is a heatmap video of vorticity (quantification of rotation), calculated using the PIV technique. The white and black arrows in the bottom panels (left and right respectively) indicate the direction and magnitude of the local velocity field of cellular motion. In the bottom panels, towards the end of the video, we see a Right-side dominance of the cellular flows, indicating a deviation from Left-Right (L-R) symmetry. All videos are synchronized and correspond to the same dataset with a total duration of 10 hours, and the timestamp represents hours and minutes.

**Movie2**: **Cellular flows in cell-division inhibited chick embryos.**
The top left panel is a brightfield imaging time-lapse video of a cell-division inhibited chick embryo, where the Primitive Streak (PS) does not form. The top right is a Flowtrace visualization video showing a Right-vortex dominated cellular flow, the cells are tagged with fluorescent labels. The bottom left is a heatmap video of cellular speeds, calculated using the Particle Image Velocimetry (PIV) technique. The bottom right is a heatmap video of vorticity (quantification of rotation), calculated using the PIV technique. The white and black arrows in the bottom panels (left and right respectively) indicate the direction and magnitude of the local velocity field of cellular motion. In the top right panel, and bottom panels, from the middle of the video until the end, we see only one Right-vortex dominated cellular flow. All videos are synchronized and correspond to the same dataset with a total duration of 10 hours, and the timestamp represents hours and minutes.

**Supplementary Figures**

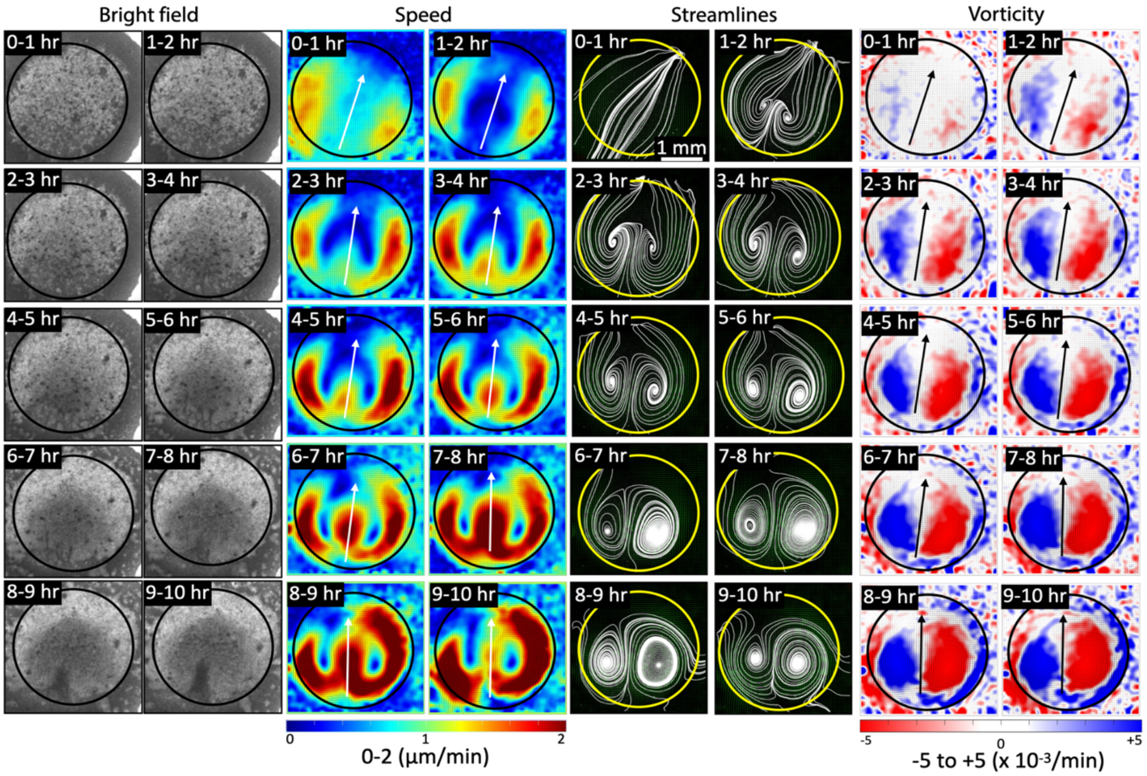


**Fig. S1. Bilateral cellular flows inside the embryonic disc.** The circular embryonic disc is shown in images of bright field, speed, streamlines and vorticity respectively. The quantification in this work only considers the region within this embryonic disc. In the vorticity calculations, the blue (L) and red (R) vortical regions are surrounded by vorticity patches of opposite colors. The patches are due to an artifact that arises because the edge of the disc has stationary extra-embryonic material, so these patches are discarded from the quantification.

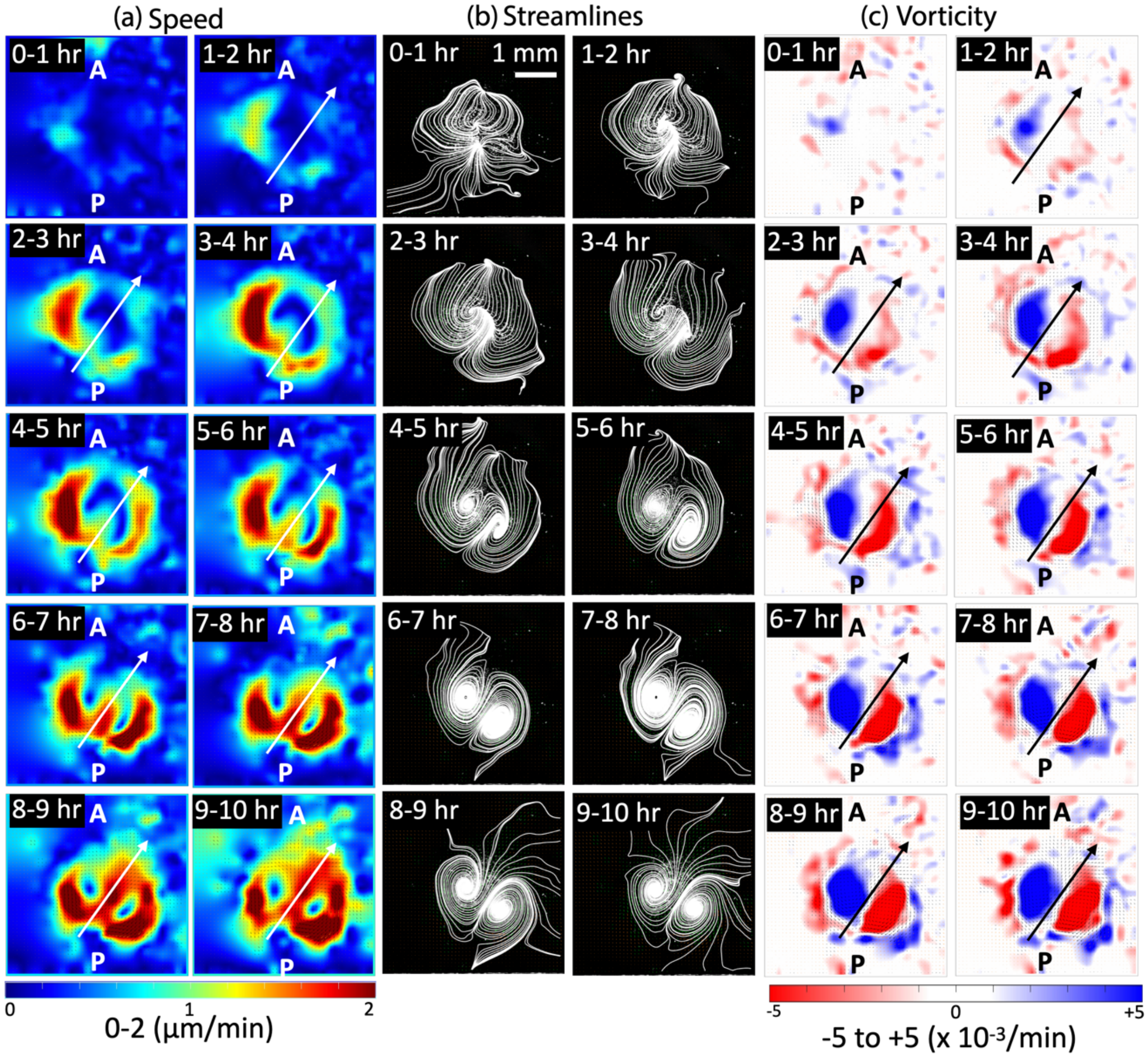


**Fig. S2. Temporal progression of bilateral flows during early chick embryo development.** Short time-averaged (1 hour) sequences of (a) speed, (b) streamlines, and (c) vorticity from PIV analysis of control sample 2 (see Table 1). The cellular speed, streamlines, and vorticity reveal that initially the cellular flows show speed transients (0-2 hours), which later stabilize into bilateral flows that are L dominated (2-4 hours). However, the bilateral flows soon regain L-R symmetry (4-5 hours) and at later stages (5 hours onwards) the cellular flows become R dominant. The arrows represent the Flow Midlines (FM) and indicate direction of flows at that location (Methods). The length scale bar shown in the first streamline panel (0-1 hour) is the same for all panels.

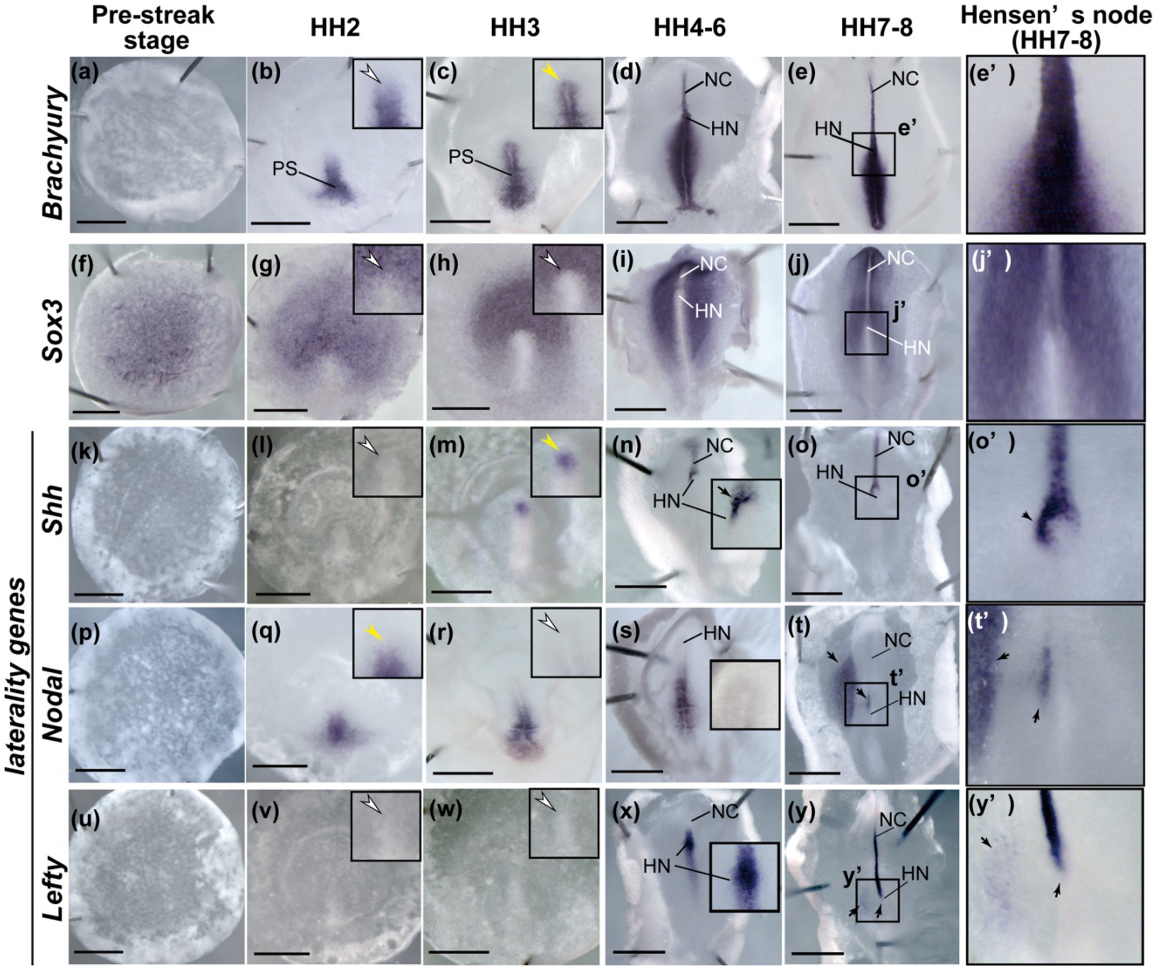


**Fig. S3. NBT/BCIP staining of WISH show that laterality genes exhibit left-right asymmetric expression pattern at Hensen's node at or after HH4.**

(a-y') Pre-streak stage to HH8 whole mount in situ hybridization of Brachyury (a-e'), Sox3 (f-j'), Shh (k-o'), Nodal (p-t'), Lefty (u-y'). The enlarged anterior tip of the PS, where the Hensen's node form at HH4, is shown in the black boxes in (b, c, g, h, l, m, q, r, v, w). White arrow heads indicate that the NBT/BCIP staining signal was not detectable. Yellow arrow heads indicate that the NBT/BCIP staining signal was detectable but the LR asymmetry in the staining signal was not evident. (e', j', o', y', y') Enlarged Hensen's node in (e), (j), (o), (t), (y), respectively. Black arrows indicate left-right asymmetric expression pattern. Scale bars, 1mm. PS; primitive streak, HN; Hensen's node, NC; notochord.

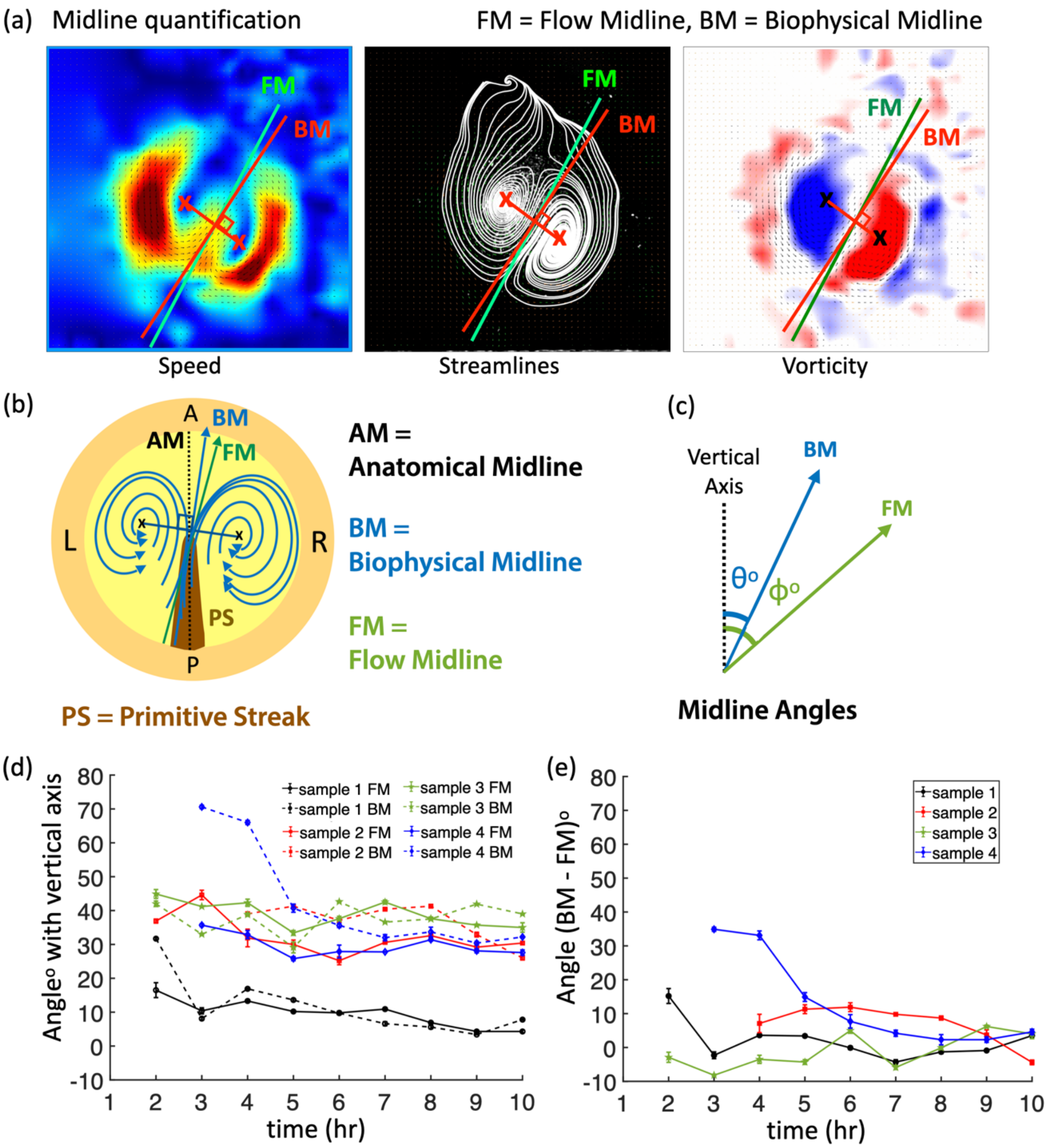


**Fig. S4. Quantification methods: Midlines.** The Biophysical Midline (BM), shown in red color, is defined as the line perpendicular to the midpoint of the line joining the centers (marked by 'X') of the two counter-rotating vortices. BM is shown for speed, streamlined vectors and vorticity in order. The Flow Midline (FM), shown in green color, is defined as the line parallel to the flow streamlines and passes through the midpoint of the line joining the vortex centers. The Anatomical Midline (AM) is the line along which the Primitive Streak (PS) forms. (b) A cartoon representation of an early-stage chicken embryo showing AM, BM, FM and two counter-rotating vortices. (c) The angles subtended by BM and FM with vertical are shown. (d) Time evolution of BM and FM angles with the vertical axis for four control samples. (e) Time evolution of angular separation between BM and FM for the control samples.

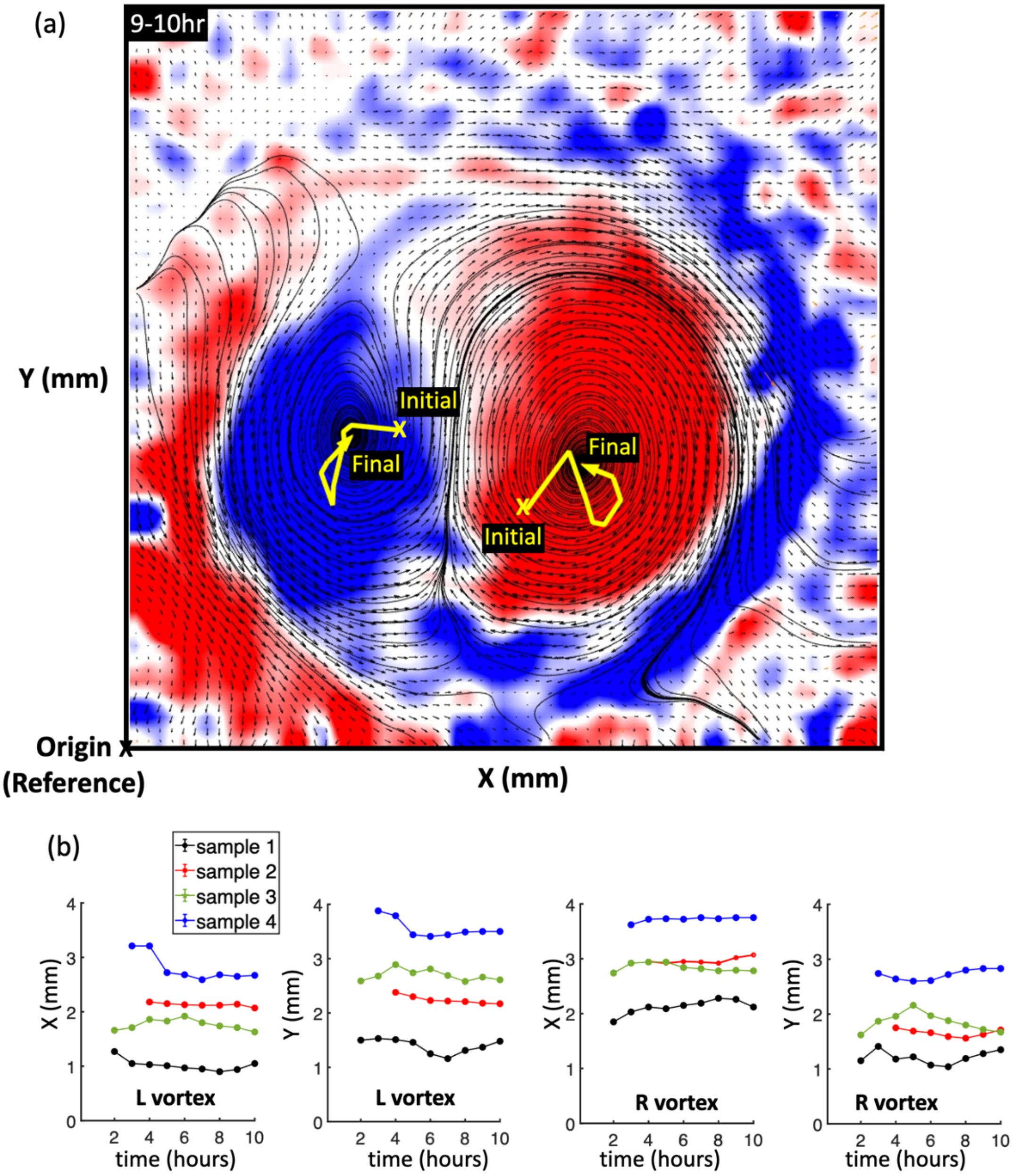


**Fig. S5. Quantification of shift in vortex centers over time.** (a) The X-Y trajectories (yellow color) of vortex centers (L and R) over time for a duration of 10 hours (the background image corresponds to 9-10 hr time window). (b) The variation of values in X and Y (in mm) are plotted over time for both the L and R vortices (N = 4 control samples). The origin (reference point) is chosen to be at the left bottom corner of the image.

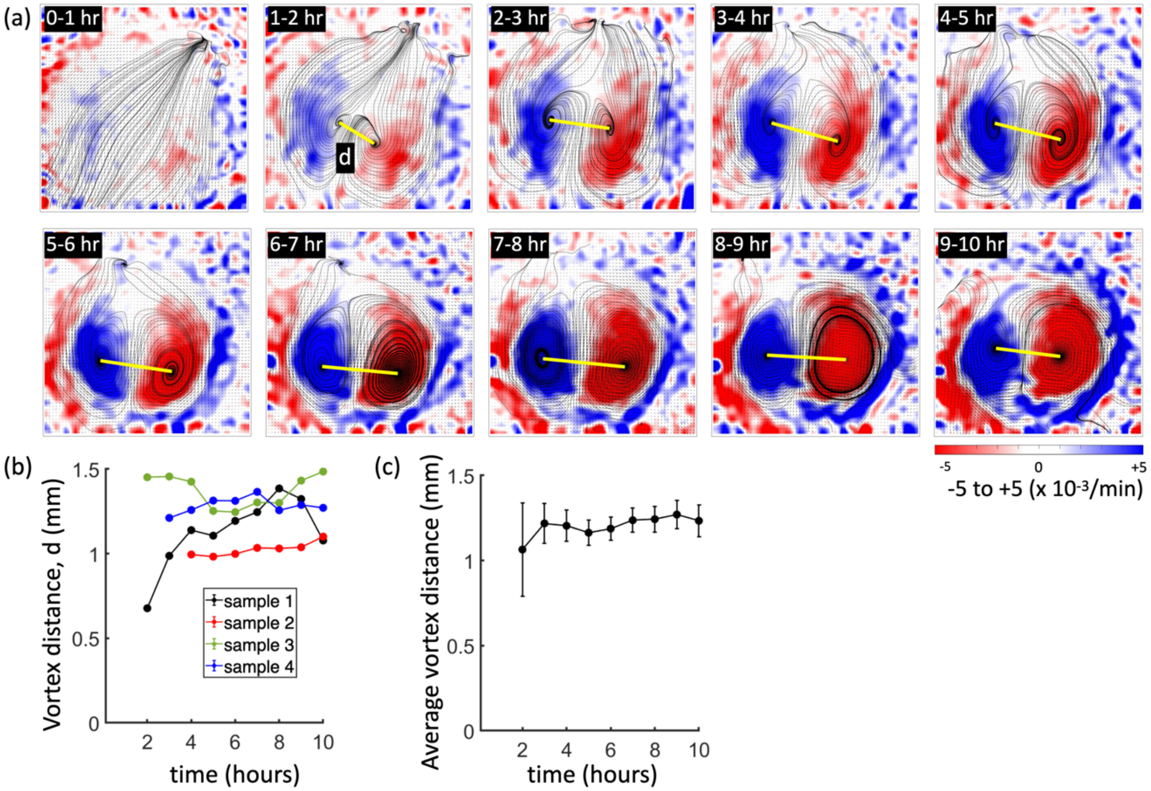


**Fig. S6. Quantification of distance between bilateral vortices:** (a) Short time-averaged (1 hour) sequence of vorticity from PIV analysis of control sample 1. The yellow line connects the two vortex centers and represents the distance between them. (b) Vortex distance, d (mm) over time (10 hours) for all control samples. (c) The average vortex distance (for N = 4 samples) over time, with the error bars representing the standard error.

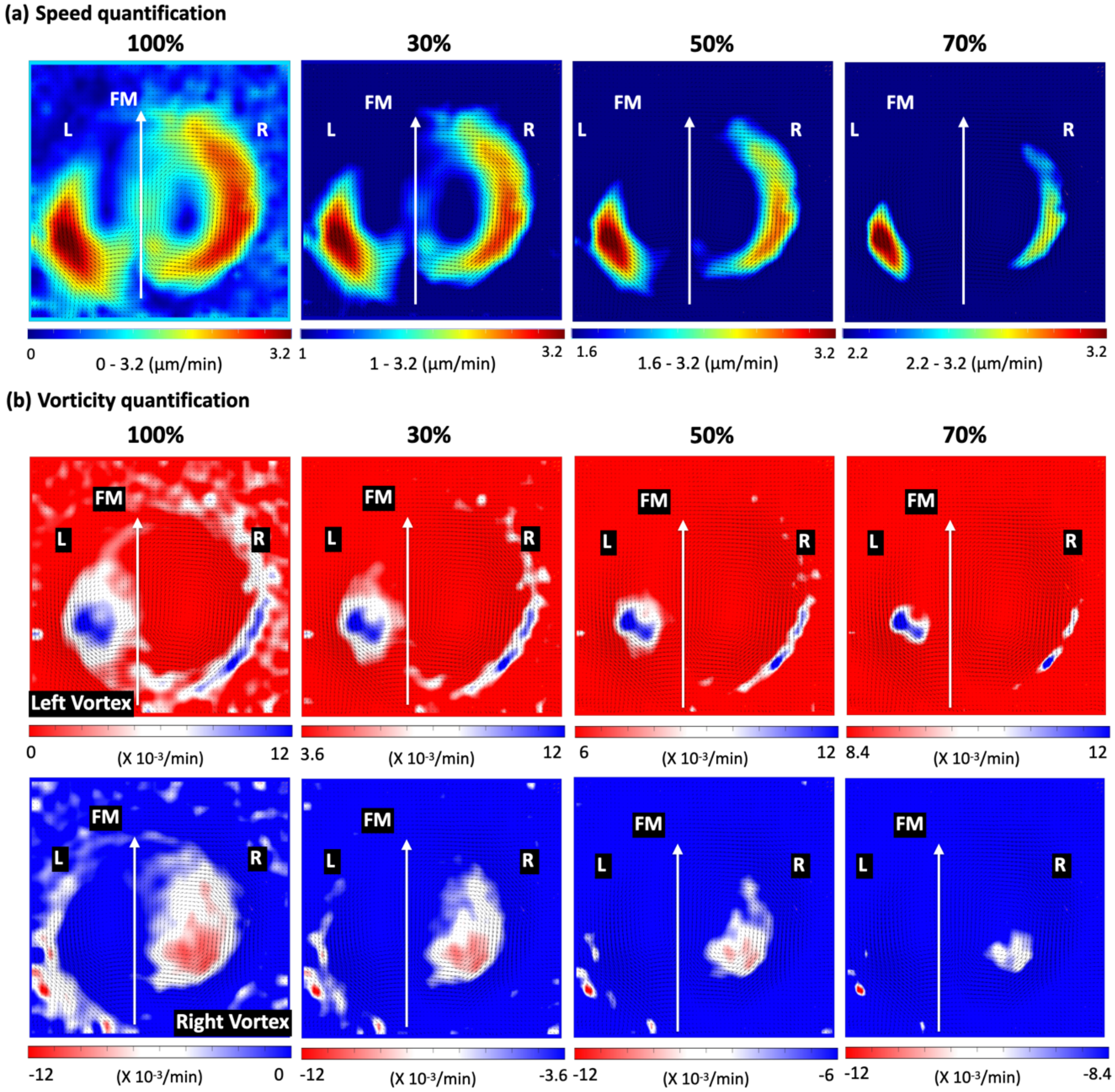


**Fig. S7. Quantification of speed and vorticity at different thresholds: (a) Speed:** The first panel shows the speed heatmap with a color bar scale between 0 - 3.2 μm/min, where 3.2 μm/min is maximum speed $V_{max}$ corresponding to the 9-10 hr time window for control sample 1 (Table S3). The next three panels show the thresholded heatmaps (30%, 50% and 70%) based on the $V_{max}$. The thresholding retains only the selected speed regions of interest, so it enables separate quantification of the thresholded area of speed in L and R regions. **(b) Vorticity:** Thresholding for the vorticity area for the two vortices (L vortex in blue and R vortex in red) based on the maximum vorticity of 12 $\times 10^{-3}$/min corresponding to the 9-10 hr time window for control sample 1 (Table S4). The three thresholds enable separate quantification of the thresholded area of vorticity in the L and R regions (blue and red respectively).

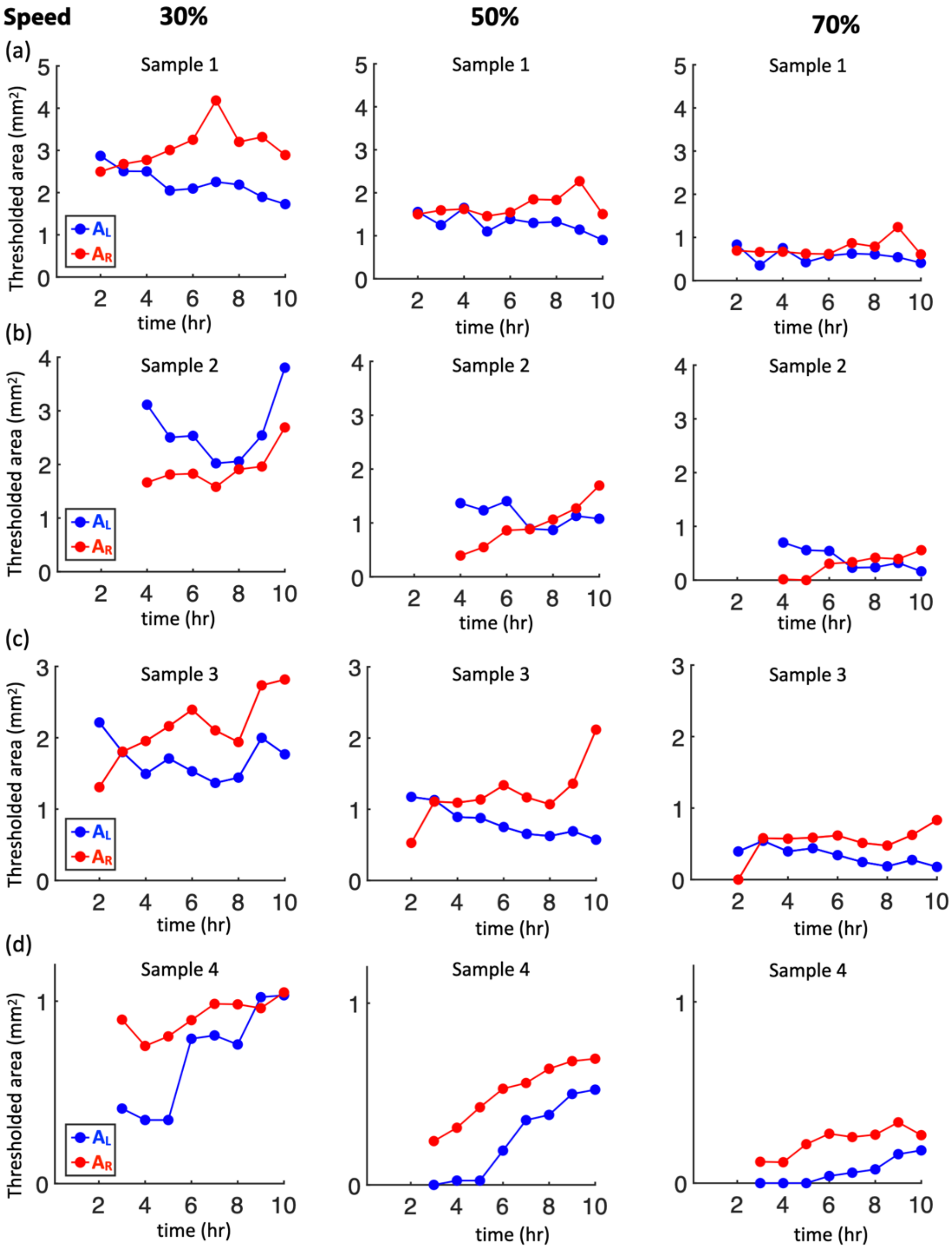


**Fig. S8. Speed quantification.** Time evolution of thresholded area of speeds in L and R regions for control samples with the three thresholds (30%, 50%, and 70%). The thresholds are based on the maximum speed $V_{max}$ (Table S3) and speeds higher than the threshold are taken to represent areas in L and R. Panels (a)-(d) represent the quantification of thresholded area of speeds (in $mm^2$) based on the thresholds for the control samples 1, 2, 3 and 4 respectively.

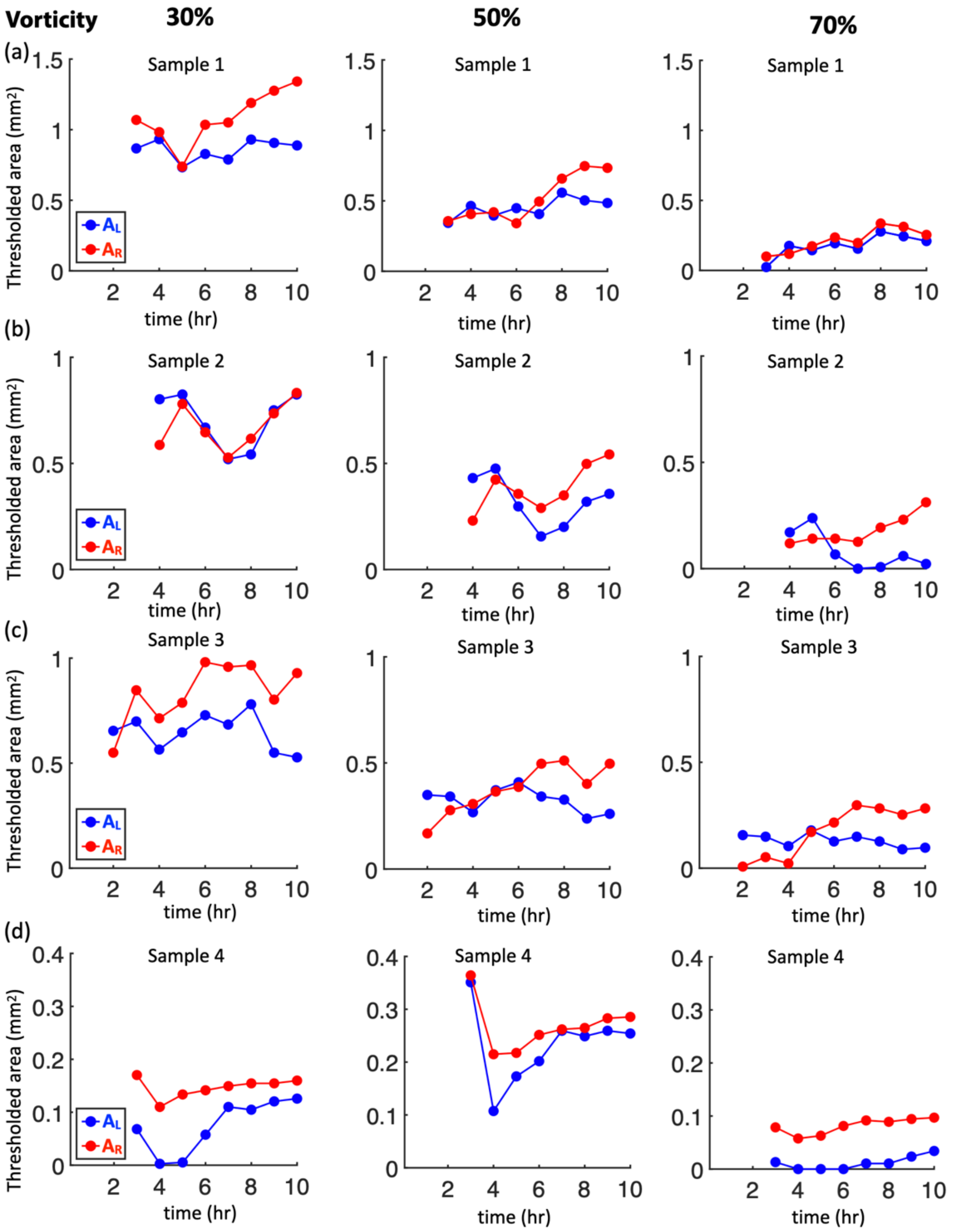


**Fig. S9. Vorticity quantification.** Time evolution of thresholded area of vorticity in L and R regions for control samples with the three thresholds (30%, 50%, and 70%). The thresholds are based on the maximum vorticity (Table S4), and vorticity values higher than the threshold are taken to represent areas in L and R. Panels (a)-(d) represent the quantification of thresholded area of vorticity (in $mm^2$) based on the thresholds for the control samples 1, 2, 3 and 4 respectively.

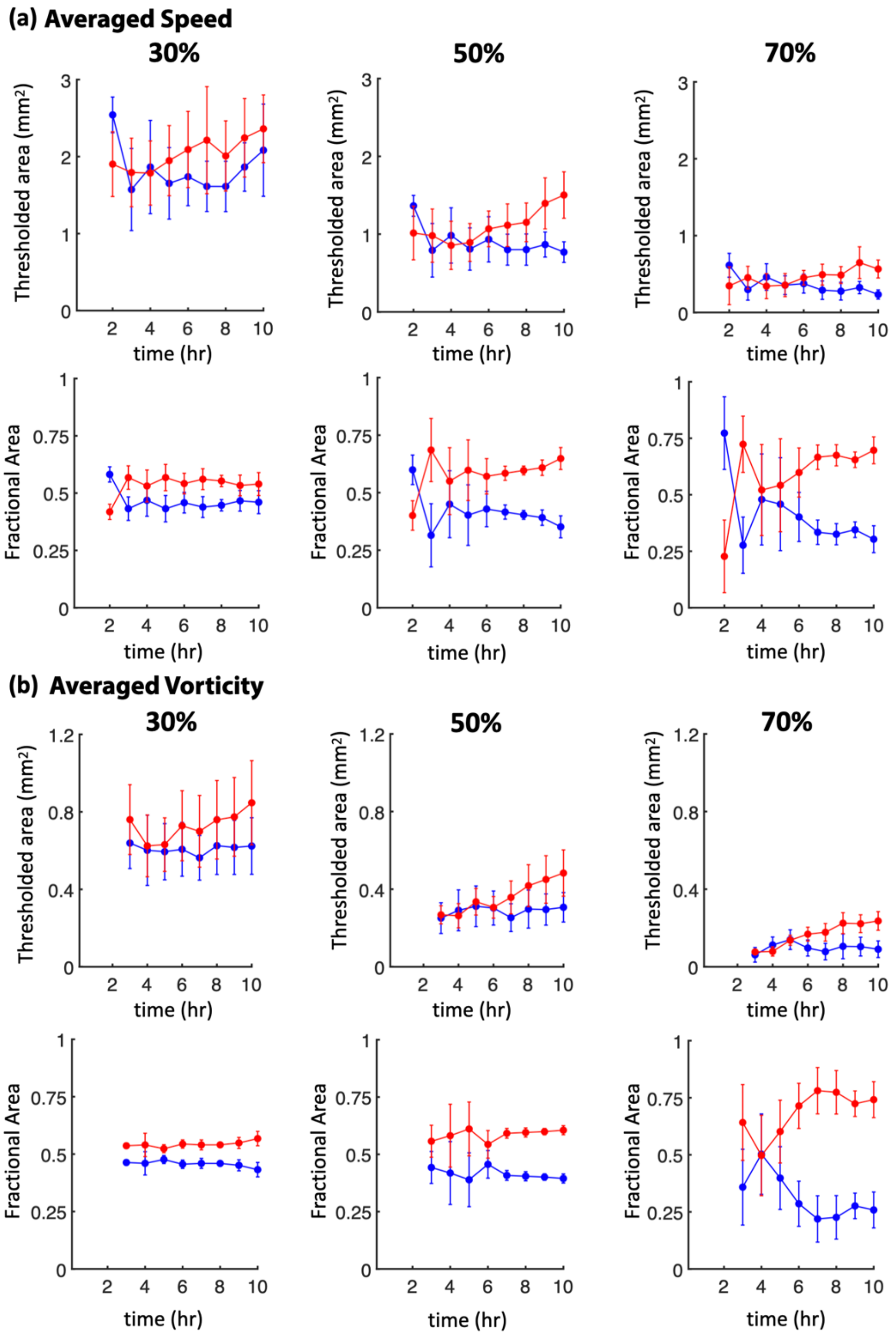


**Fig. S10. Averaged speed and vorticity quantification over time. (a) Speed:** The first row shows the time evolution of the averaged mean (N = 4 samples) of thresholded area of speeds in L and R regions for the respective thresholds. Lower thresholds (30%) allow more values to contribute to larger areas, whereas a higher (stricter) threshold (70%) will retain fewer number of values of high speeds, and hence result in lower area values. The second row represents the time evolution of the fractional speed area in the L and R regions, calculated as ($A_L/A_L+A_R$) and ($A_R/A_L+A_R$) for different thresholds. **(b) Vorticity:** The first row shows the thresholded area of vorticity (N = 4 samples) in the L and R regions for the three thresholds. The second row shows the fractional vorticity area in the L and R sides ($A_L/A_L+A_R$) and ($A_R/A_L+A_R$). All error bars shown represent the standard error.

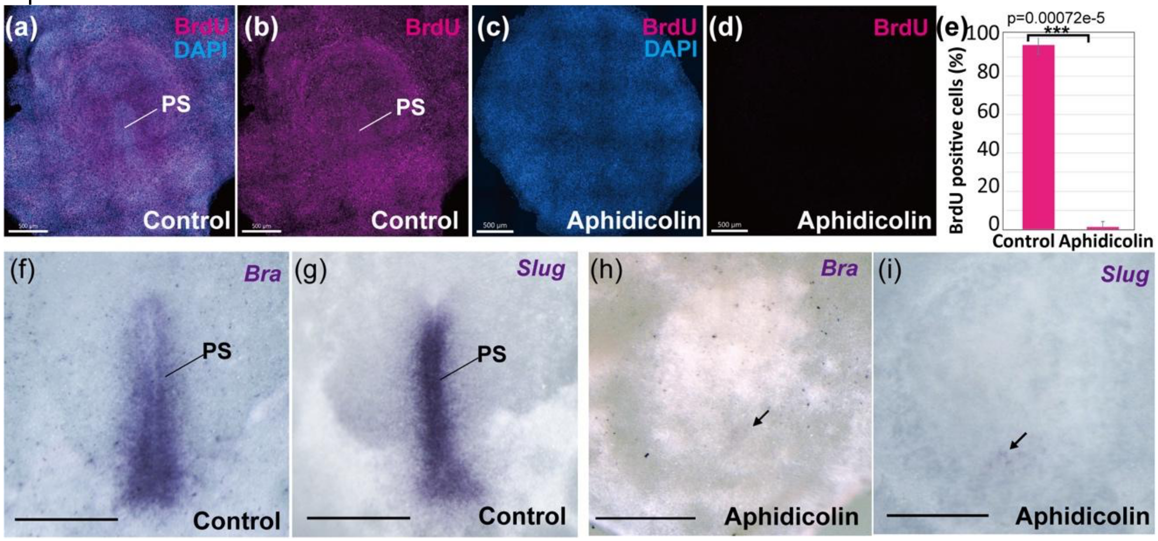


**Figure S11. Mitotic arrest diminished midline structure.** (a-d) BrdU-incorporation in control (a, b) and aphidicolin-treated embryos (c, d). (e) BrdU-positive cells (%) in control and aphidicolin-treated embryos (control 96.2±4.8% vs aphidicolin 1.42±2.8%, p=0.00072e-5, N=5 for each, Student's T-test). (f-i) Stage HH3 whole mount in situ hybridization of Brachyury (Bra; f, h) and Slug (g, i). PS, the primitive streak. N = 4 for each. Scale bars, 500 µm.

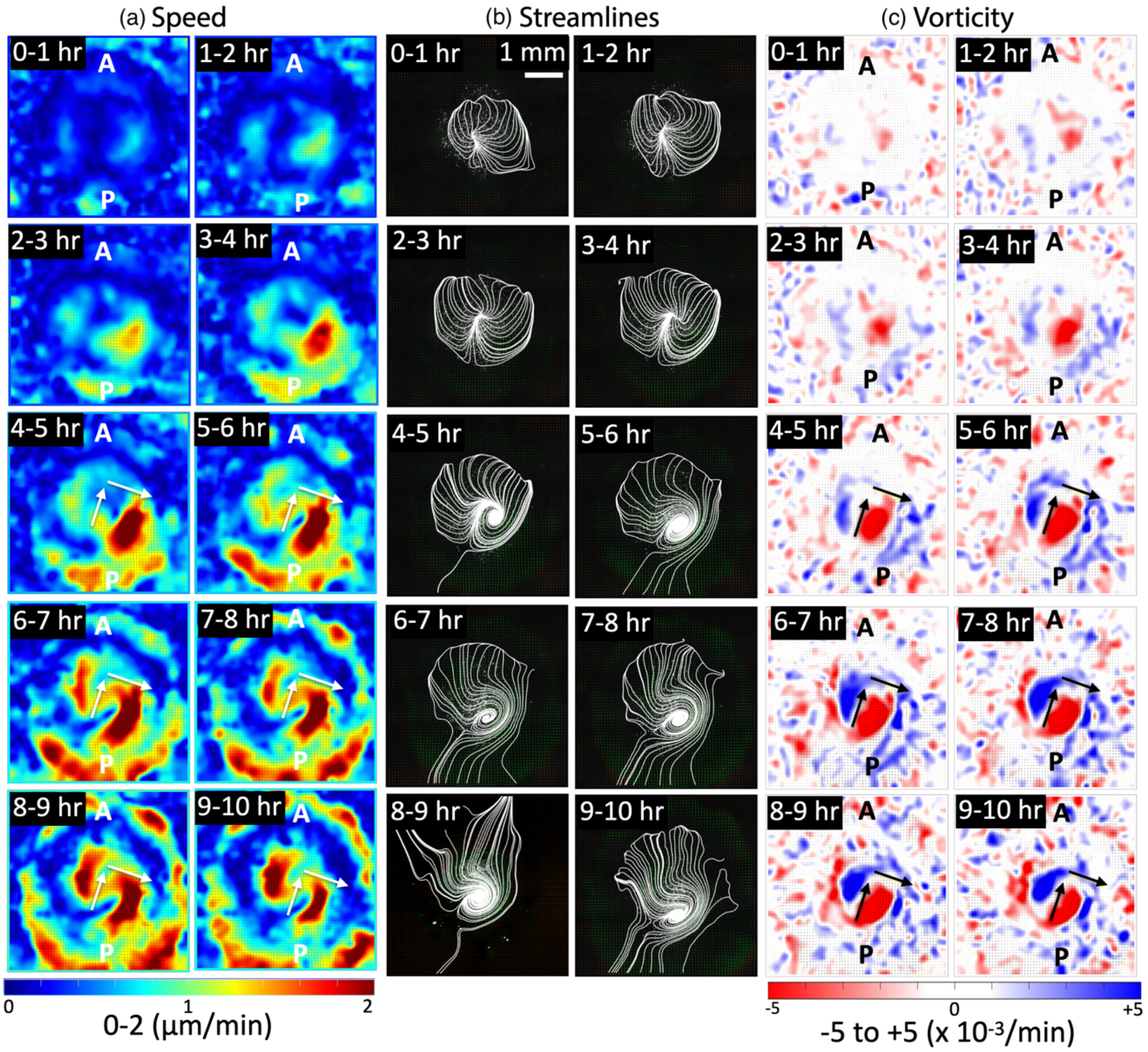


**Fig. S12. Temporal progression of bilateral flows in cell-division inhibited chick embryos.** Short time-averaged (1 hour) sequences of (a) speed, (b) streamlines, and (c) vorticity from PIV analysis of aphidicolin sample 2 (see Table 1). The cellular speed, streamlines, and vorticity reveal that the initial cellular flow transients (0-2 hours) do not stabilize to form bilateral flows (2 hours onwards). However, the cellular flows consist of only one vortex that is R dominated (2 hours onwards until the end). Although the vorticity calculations pick-up features of counterclockwise rotational flows in blue color (4 hours onwards), it is important to note that there is only one well-defined R vortex in this dataset. The short arrows serve as a guide to indicate direction of flows at that location, but do not represent midlines (Methods). The length scale bar shown in the first streamline panel (0-1 hour) is the same for all panels.

**Table S1.** PIV analysis settings for different datasets.

| Sample Number | Window size (px) pass 1 | Window width (px) pass 2 |
|---|---|---|
| Control Sample 1 | 128 x 128 | 64 x 64 |
| Control Sample 2 | 64 x 64 | 32 x 32 |
| Control Sample 3 | 64 x 64 | 32 x 32 |
| Control Sample 4 | 128 x 128 | 64 x 64 |
| Aphidicolin Sample 1 | 128 x 128 | 64 x 64 |
| Aphidicolin Sample 2 | 128 x 128 | 64 x 64 |
| Aphidicolin Sample 3 | 128 x 128 | 64 x 64 |
| Aphidicolin Sample 4 | 128 x 128 | 64 x 64 |

**Table S2.** Anatomical Midline (AM) angle in control embryos.

| Sample number | Angle with vertical (AM) in degrees |
|---|---|
| Control Sample 1 | $6 \pm 0.2$ |
| Control Sample 2 | $37 \pm 1.4$ |
| Control Sample 3 | $31.6 \pm 1.0$ |
| Control Sample 4 | $31.0 \pm 0.9$ |

**Table S3**: Quantification of speed thresholds (30%, 50%, 70%) based on maximum speed ($V_{max}$) at respective hourly time windows in control embryos.

Sample 1

| Time | $V_{max}$ (⌠m/min) | 30% | 50% | 70% |
|---|---|---|---|---|
| 0-1 hr | - | - | - | - |
| 1-2 hr | 1.4 | 0.42 | 0.70 | 0.98 |
| 2-3 hr | 1.8 | 0.54 | 0.90 | 1.26 |
| 3-4 hr | 1.8 | 0.54 | 0.90 | 1.26 |
| 4-5 hr | 2.2 | 0.66 | 1.10 | 1.54 |
| 5-6 hr | 2.1 | 0.63 | 1.00 | 1.47 |
| 6-7 hr | 2.2 | 0.66 | 1.10 | 1.54 |
| 7-8 hr | 2.6 | 0.78 | 1.30 | 1.82 |
| 8-9 hr | 2.6 | 0.78 | 1.30 | 1.82 |
| 9-10 hr | 3.2 | 0.96 | 1.60 | 2.24 |

Sample 2

| Time | $V_{max}$ (⌠m/min) | 30% | 50% | 70% |
|---|---|---|---|---|
| 0-1 hr | - | - | - | - |
| 1-2 hr | - | - | - | - |
| 2-3 hr | - | - | - | - |
| 3-4 hr | 2.3 | 0.69 | 1.15 | 1.61 |
| 4-5 hr | 2.5 | 0.75 | 1.25 | 1.75 |
| 5-6 hr | 2.3 | 0.69 | 1.15 | 1.61 |
| 6-7 hr | 2.7 | 0.81 | 1.35 | 1.89 |
| 7-8 hr | 2.8 | 0.84 | 1.40 | 1.96 |
| 8-9 hr | 2.8 | 0.84 | 1.40 | 1.96 |
| 9-10 hr | 2.8 | 0.84 | 1.40 | 1.96 |

Sample 3

| Time | $V_{max}$ (⌠m/min) | 30% | 50% | 70% |
|---|---|---|---|---|
| 0-1 hr | - | - | - | - |
| 1-2 hr | 1.4 | 0.42 | 0.70 | 0.98 |
| 2-3 hr | 1.6 | 0.48 | 0.80 | 1.12 |
| 3-4 hr | 1.8 | 0.54 | 0.90 | 1.26 |
| 4-5 hr | 2.0 | 0.60 | 1.00 | 1.40 |
| 5-6 hr | 1.8 | 0.54 | 0.90 | 1.26 |
| 6-7 hr | 1.6 | 0.48 | 0.80 | 1.12 |
| 7-8 hr | 1.7 | 0.51 | 0.85 | 1.19 |
| 8-9 hr | 1.7 | 0.51 | 0.85 | 1.19 |
| 9-10 hr | 1.8 | 0.54 | 0.90 | 1.26 |

Sample 4

| Time | $V_{max}$ (⌠m/min) | 30% | 50% | 70% |
|---|---|---|---|---|
| 0-1 hr | - | - | - | - |
| 1-2 hr | - | - | - | - |
| 2-3 hr | 1.2 | 0.36 | 0.60 | 0.84 |
| 3-4 hr | 1.4 | 0.42 | 0.70 | 0.98 |
| 4-5 hr | 1.8 | 0.54 | 0.90 | 1.26 |
| 5-6 hr | 2.2 | 0.66 | 1.10 | 1.54 |
| 6-7 hr | 2.4 | 0.72 | 1.12 | 1.68 |
| 7-8 hr | 2.4 | 0.72 | 1.12 | 1.68 |
| 8-9 hr | 2.4 | 0.72 | 1.12 | 1.68 |
| 9-10 hr | 2.4 | 0.72 | 1.12 | 1.68 |

**Table S4**: Vorticity thresholds based on maximum vorticity ($Vorticity_{max}$) at respective time points for four control samples.

Sample 1

| Time | $Vorticity_{max}$ ( $x10^{-3}$/min) | 30% | 50% | 70% |
|---|---|---|---|---|
| 0-1 hr | - | - | - | - |
| 1-2 hr | - | - | - | - |
| 2-3 hr | 8 | 2.40 | 4 | 5.56 |
| 3-4 hr | 8.6 | 2.58 | 4.43 | 6.00 |
| 4-5 hr | 10 | 3 | 5 | 7.00 |
| 5-6 hr | 10 | 3 | 5 | 7.00 |
| 6-7 hr | 12 | 3.6 | 6 | 8.40 |
| 7-8 hr | 12 | 3.6 | 6 | 8.40 |
| 8-9 hr | 12 | 3.6 | 6 | 8.4 |
| 9-10 hr | 12 | 3.6 | 6 | 8.4 |

Sample 2

| Time | $Vorticity_{max}$ ( $x10^{-3}$/min) | 30% | 50% | 70% |
|---|---|---|---|---|
| 0-1 hr | - | - | - | - |
| 1-2 hr | - | - | - | - |
| 2-3 hr | - | - | - | - |
| 3-4 hr | 9 | 2.70 | 4.50 | 6.30 |
| 4-5 hr | 9 | 2.70 | 4.50 | 6.30 |
| 5-6 hr | 12 | 3.6 | 6.00 | 8.40 |
| 6-7 hr | 16 | 4.80 | 8.00 | 11.2 |
| 7-8 hr | 16 | 4.80 | 8.00 | 11.2 |
| 8-9 hr | 14 | 4.20 | 7.00 | 9.80 |
| 9-10 hr | 13 | 3.9 | 6.5 | 9.1 |

Sample 3

| Time | $Vorticity_{max}$ ( $x10^{-3}$/min) | 30% | 50% | 70% |
|---|---|---|---|---|
| 0-1 hr | - | - | - | - |
| 1-2 hr | 6 | 1.8 | 3 | 4.2 |
| 2-3 hr | 7 | 2.1 | 3.5 | 4.9 |
| 3-4 hr | 8 | 2.40 | 4 | 5.56 |
| 4-5 hr | 8 | 2.40 | 4 | 5.56 |
| 5-6 hr | 7 | 2.1 | 3.5 | 4.9 |
| 6-7 hr | 6 | 1.8 | 3 | 4.2 |
| 7-8 hr | 6.5 | 1.95 | 3.25 | 4.55 |
| 8-9 hr | 8.5 | 2.55 | 4.25 | 5.95 |
| 9-10 hr | 9 | 2.70 | 4.50 | 6.30 |

Sample 4

| Time | $Vorticity_{max}$ ( $x10^{-3}$/min) | 30% | 50% | 70% |
|---|---|---|---|---|
| 0-1 hr | - | - | - | - |
| 1-2 hr | - | - | - | - |
| 2-3 hr | 3.4 | 1 | 1.7 | 2.38 |
| 3-4 hr | 5.8 | 1.74 | 2.9 | 4 |
| 4-5 hr | 7.9 | 2.37 | 3.95 | 5.53 |
| 5-6 hr | 10 | 3 | 5 | 7.00 |
| 6-7 hr | 10 | 3 | 5 | 7.00 |
| 7-8 hr | 10 | 3 | 5 | 7.00 |
| 8-9 hr | 10 | 3 | 5 | 7.00 |
| 9-10 hr | 10 | 3 | 5 | 7.00 |

**Table S5**: Wilcoxon test p-value for L-R fractional area of speed over time.

| Time | 30% | 50% | 70% |
|---|---|---|---|
| 0-1 hr | - | - | - |
| 1-2 hr | 0.24 | 0.24 | 0.24 |
| 2-3 hr | 0.08 | 0.19 | 0.08 |
| 3-4 hr | 0.47 | 0.66 | 0.66 |
| 4-5 hr | 0.19 | 0.38 | 0.66 |
| 5-6 hr | 0.19 | 0.19 | 0.19 |
| 6-7 hr | 0.19 | 0.06 | 0.03 |
| 7-8 hr | 0.06 | 0.03 | 0.03 |
| 8-9 hr | 0.31 | 0.03 | 0.03 |
| 9-10 hr | 0.19 | 0.03 | 0.03 |

**Table S6**: Wilcoxon test p-value for L-R fractional area of vorticity over time.

| Time | 30% | 50% | 70% |
|---|---|---|---|
| 0-1 hr | - | - | - |
| 1-2 hr | - | - | - |
| 2-3 hr | 0.08 | 0.66 | 0.19 |
| 3-4 hr | 0.47 | 0.66 | 0.88 |
| 4-5 hr | 0.19 | 0.66 | 0.66 |
| 5-6 hr | 0.06 | 0.66 | 0.03 |
| 6-7 hr | 0.03 | 0.03 | 0.03 |
| 7-8 hr | 0.03 | 0.03 | 0.03 |
| 8-9 hr | 0.06 | 0.03 | 0.03 |
| 9-10 hr | 0.03 | 0.03 | 0.03 |
